# Dysfunction of the Polycomb protein RYBP and of 5-methylcytosine oxidases leads to widespread CpG island hypermethylation and cell transformation

**DOI:** 10.1101/2022.07.26.501603

**Authors:** Wei Cui, Zhijun Huang, Seung-Gi Jin, Jennifer Johnson, Galen Hostetter, Gerd P. Pfeifer

**Author notes:** Correspondence: Gerd P. Pfeifer Department of Epigenetics Van Andel Institute Grand Rapids, MI 49525 1-616-234-5398. These authors contributed equally (Wei Cui, Zhijun Huang).

## Abstract

DNA hypermethylation is a hallmark of cancer and predominantly affects CpG island regions. Although this phenomenon was first described more than three decades ago, its mechanisms have remained unknown. Since CpG island hypermethylation is strongly biased towards Polycomb target genes, we reasoned that dysfunction of Polycomb repression complexes (PRCs) may underlie CpG island hypermethylation. We observed that a few genes coding for components of the PRC1 complex are downregulated in many cancer types. We focused on RYBP, a key activator of variant PRC1 complexes responsible for H2AK119 monoubiquitylation. We inactivated RYBP in nontumorigenic bronchial epithelial cells and observed a limited extent of DNA hypermethylation. Considering that tumors are deficient in 5-methylcytosine oxidase (TET protein) function as documented by substantially reduced levels of 5-hydroxymethylcytosine in all solid tumors, we then inactivated TET1, TET2, and TET3 in bronchial cells, individually and in combination. Using quadruple knockouts of RYBP and all three TET proteins, we observed widespread hypermethylation of H2AK119Ub1-marked CpG islands affecting almost 4,000 target genes. This hypermethylation closely mirrored the DNA hypermethylation landscape observed in human lung tumors. These cells showed aberrant methylation and dysregulation of several cancer-relevant pathways including cell cycle control genes, defects in the Hippo pathway and overexpression of AP-1 transcription factor genes. As a result, the quadruple knockout bronchial cells acquired properties of a transformed phenotype, including efficient growth in soft agar and formation of squamous cell carcinomas in immune-compromised mice. Our data provide a long-sought mechanism for DNA hypermethylation in cancer and explain how such hypermethylation leads to cell transformation. Cancer formation, therefore, is achievable by misregulation of two epigenetic pathways without introduction of cancer driver mutations.

---

Epigenetic reprogramming is a hallmark of cancer (Hanahan, 2022). Changes of the epigenome in tumors (Baylin and Jones, 2016) include altered DNA methylation patterns and aberrant nucleosome modifications. Cancer cells have long been known to harbor extensive genome-wide DNA hypomethylation relative to the corresponding normal cells and show focal DNA hypermethylation at CpG islands (CGIs) (Baylin et al., 1986; Feinberg and Vogelstein, 1983; Gama-Sosa et al., 1983). Although the consequences of DNA hypomethylation for tumorigenesis are less clear, DNA hypermethylation correlates with silencing of several tumor suppressor genes, for example genes involved in cell cycle control, anti-growth signaling pathways, or DNA repair (Jones and Laird, 1999; Lahtz and Pfeifer, 2011; Merlo et al., 1995). DNA hypermethylation of CpG islands is widespread, affecting a thousand or more target genes in individual tumors and is seen across most cancer types. This event does not occur randomly but is strongly targeted to genes encoding developmental transcription factors and signaling molecules that are regulated by the Polycomb complex (Ohm et al., 2007; Rauch et al., 2006; Rauch et al., 2007; Schlesinger et al., 2007; Widschwendter et al., 2007).

DNA methylation patterns in normal and malignant cells are established by DNA methyltransferases (DNMTs), which operate preferentially at 5’CpG sequences and produce 5- methylcytosine (5mC). DNMT1 maintains methylation patterns owing to its preferential activity on hemimethylated CpG sites formed shortly after DNA replication, and DNMT3A and DNMT3B can methylate unmethylated sites *de novo* (Edwards et al., 2017). DNA methylation patterns can be reversed to an unmethylated state by enzymatic activities that covert the methyl group of 5- methylcytosine to oxidized forms creating 5-hydroxymethyl-, 5-formyl-, and 5-carboxylcytosine followed by base excision repair of the latter two bases. These reactions are carried out by a small family of 5-methylcytosine oxidases, the TET proteins (TET1, TET2, and TET3) (An et al., 2017; Pfeifer et al., 2013; Tahiliani et al., 2009; Wu and Zhang, 2017). DNMT and TET proteins are found mutated in some human tumors, but this process primarily affects DNMT3A and TET2 in hematological malignancies (Abdel-Wahab et al., 2009; Yang et al., 2015). On a more global scale, the 5-methylcytosine oxidation process is defective in all human solid tumors analyzed, independent of specific genomic mutations in genes encoding TET proteins or genes critical in the biochemical pathways that support 5mC oxidation (Jin et al., 2011).

Although DNA methylation patterns in human cancer have been extensively characterized, we still have no good understanding of how these methylation changes arise mechanistically. We hypothesized that the specificity of DNA hypermethylation for Polycomb target genes is based on a diminished function of Polycomb and is accompanied by defective 5mC oxidation. In a model system using human bronchial epithelial cells, we have inactivated a critical component of PRC1 complexes, RYBP, and the TET proteins and analyzed the consequences of these deficiencies on genome-wide DNA methylation and cellular phenotype.

## Inactivation of RYBP in bronchial cells

We first analyzed The Cancer Genome Atlas (TCGA) database and examined the expression of genes coding for subunits of Polycomb repression complexes (PRC1 and PRC2). Among over 50 genes analyzed for PRC1 and PRC2, we found that most Polycomb genes were upregulated in tumors across a wide spectrum of malignancies. The only downregulated genes in lung cancer and other tumors were *EZH1* for PRC2 and *RYBP*, *PCGF5*, *CBX6* (in lung adenocarcinomas), and *CBX7* for PRC1 (Fig. 1a; Extended Data Fig. 1a,b). *EZH1* downregulation is accompanied by a strong upregulation of *EZH2* as seen in many cancer types (TCGA database). We decided to focus on RYBP because this protein is present in most if not all variant (also called noncanonical) PRC1 complexes where it is mutually exclusive with CBX proteins (Gao et al., 2012; Piunti and Shilatifard, 2021; Tavares et al., 2012). Variant PRC1 complexes are responsible for the largest fraction of total H2AK119 monoubiquitylation in cells (Blackledge and Klose, 2021). RYBP is a strong activator of the RING1B ubiquitin ligase activity of PRC1 that produces H2AK119Ub1 (Morey et al., 2013; Rose et al., 2016) and provides a positive feedback loop for focused PRC1 activities through its binding to H2AK119ub1 (Blackledge and Klose, 2021; Zhao et al., 2020). RYBP is downregulated in lung adeno- and squamous cell carcinomas (Fig. 1a) but also across many other types of solid tumors (Extended Data Fig. 1a). Since the focus of our research has been on DNA methylation in lung cancers (Rauch et al., 2007; Rauch et al., 2012; Rauch et al., 2008), we proceeded to inactivate RYBP in human bronchial epithelial cells. We used the telomerase- and CDK4-immortalized but nontumorigenic bronchial cell line HBEC3-KT (Ramirez et al., 2004; Sato et al., 2006; Vaz et al., 2017) and employed CRISPR-Cas9 technology to target the RYBP gene.

**Fig. 1.**
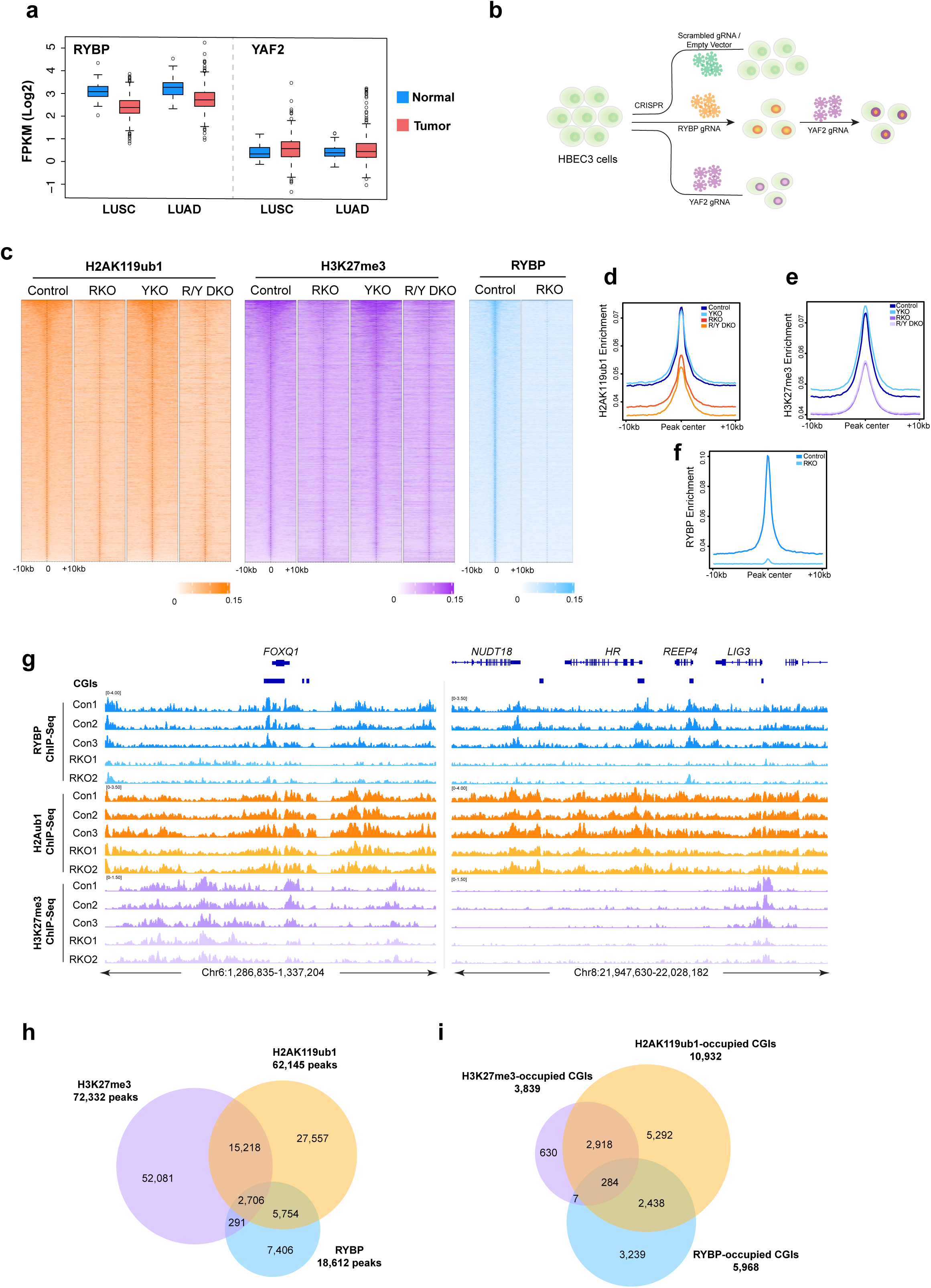
Polycomb marks in control bronchial cells and in cells lacking RYBP. **a.** mRNA expression of *RYBP* and *YAF2* in human lung adenocarcinomas (LUAD, n=537; normal solid tissue, n=59) and lung squamous cell carcinoma (LUSC, n=502; normal solid tissue, n=49) in the TCGA dataset. **b.** Strategic overview to generate knockout clones that were infected with gRNA lentivirus. **c.** Heatmap of H2AK119ub1 and H3K27me3 peaks (±10 kb) in control, RYBP knockout (RKO), YAF2 knockout (YKO) and RYBP/YAF2 double knockout (R/Y DKO) and heatmap of RYBP peaks (±10 kb) in control and RKO cells. The ChIP-Seq signal is ranked from highest to lowest for each mark in control cells. **d.** ChIP-seq cumulative enrichment deposition centered at peak summit of H2AK119ub1 in control, RKO, YKO and R/Y DKO cells. **e.** ChIP-seq cumulative enrichment deposition centered at peak summit of H3K27me3 in control, RKO, YKO and R/Y DKO cells. **f.** ChIP-seq cumulative enrichment deposition centered at peak summit of RYBP in control and RKO cells. **g.** IGV browser view of the *FOXQ1* gene, and a cluster of genes, *NUDT18*, *HR*, *REEP4*, *LIG3* for RYBP, H2AK119ub1 and HEK27me3 ChIP-seq data acquired from control and RKO cells. Three independent clones in control cells and two independent clones in RKO cells are presented**. h.** Venn diagram showing overlap of H2AK119ub1, H3K27me3 and RYBP peaks in HBEC3 control cells. **i.** Venn diagram showing overlap of CGIs targeted by H2AK119ub1, H3K27me3 and RYBP peaks in HBEC3 control cells.

Creating frameshift mutations (Extended Data Fig. 1c; Supplementary Table 1), we generated several RYBP-deficient clones, along with control clones (Fig. 1b; Extended Data Fig. 1c,d). Since RYBP has a paralogue in the human genome, YAF2, we also inactivated YAF2 (Extended Data Fig. 1e), alone or in combination with RYBP, creating a double knockout (Extended Data Fig. 1f). YAF2 is not differentially expressed in lung tumors relative to normal lung (Fig. 1a). We next assessed the levels of the PRC1 Polycomb mark, H2AK119Ub1, and the PRC2 Polycomb mark, H3K27me3, considering the extensive crosstalk between PRC1 and PRC2 activities (Blackledge and Klose, 2021; Piunti and Shilatifard, 2021). Western blotting to assess global levels of these marks showed a reduction of H2AK119Ub1 and to a lesser extent of H3K27me3 in the RYBP inactivated cells or the RYBP/YAF2 double knockouts (DKOs) but not in the single YAF2 knockout (Extended Data Fig. 1g-i).

We then performed chromatin immunoprecipitation sequencing (ChIP-seq) of the two Polycomb marks and of RYBP protein in control and RYBP/YAF2 targeted cells (Fig. 1c-i) and determined peak distributions (Supplementary Table 2, Excel file). This data shows an almost 50% reduction of H2AK119ub1 signal in cells lacking RYBP (Fig. 1c,d) and a nearly complete loss of RYBP peaks in RYBP-deleted cells (Fig. 1c,f). Loss of H3K27me3 was also substantial (Fig. 1c,e). Examples of ChIP-seq data for H2AK119ub1 across two gene loci is shown in Figure 1g. To analyze the overlap between H2AK119ub1, H3K27me3, and RYBP peaks, we created Venn Diagrams (Fig. 1h,i). There was a partial overlap between the two histone marks but less than 10% of all peaks were common to all three mapped parameters. This data shows that PRC1 and PRC2 activities do not always coincide in these somatic cells. It is also of interest that PCR1 complexes containing RYBP have in some studies been associated with active rather than repressed genes (van den Boom et al., 2016), which may explain some of the less than complete overlap. However, considering CpG islands (CGIs), about 40% of the RYBP-targeted CpG islands also carried the H2AK119ub1 mark (Fig. 1i). Of note, comparison with data for H3K4me3 mapped in bronchial epithelial cells by ENCODE revealed that the H3K27me3-associated CGIs did not carry H3K4me3 in somatic lung epithelial cells, in other words, they were not ‘bivalent’ CGIs. Focusing on CGIs marked by H2AK119ub1 alone or in combination with RYBP, we performed gene ontology analysis. These CGIs were associated with genes involved in developmental and differentiation processes (Extended Data Fig. 2a,b). In addition, KEGG pathway analysis revealed that the RYBP/H2AK119ub1 co-occupied CGIs are linked to genes involved in several cancer-relevant pathways including, for example AKT signaling and the Hippo pathway (Extended Data Fig. 2c).

Next, we performed RNA-seq in the RYBP, YAF2, and RYBP/YAF2 double knockout cells (Extended Data Fig. 2d-h). Somewhat surprisingly, YAF2 single knockout cells did not show any differentially expressed genes that reached genome-wide statistical significance. RYBP single and RYBP/YAF2 double knockouts had about 1,000 upregulated genes, which was >3-times more than the number of downregulated genes, consistent with a mostly repressive function of RYBP (Extended Data Fig. 2d-f; Supplementary Table 3, Excel file). Upregulated genes showed a loss of the H2AK119ub1 and H3K27me3 marks near the TSS, and downregulated genes showed a small increase of these repressive marks (Extended Data Fig. 2e,f). Gene ontology analysis revealed that the differentially expressed genes after loss of RYBP fell into several cancer-relevant pathways including cell migration, proliferation, epithelial to mesenchymal transition, and signal transduction (Extended Data Fig. 2g). Upregulated genes included several transcription factors, for example *ERG1*, *ERG2*, *NR4A1* and *ATF3* (Extended Data Fig. 2h).

We then determined if the loss of RYBP and PRC1/2 histone marks leads to a change in DNA methylation patterns. We analyzed three control clones and three independently derived RYBP or YAF2 knockout clones by whole genome bisulfite sequencing (WGBS), a technique that comprehensively interrogates all CpG sites in the human genome that have sufficient read coverage. DMR-seq analysis (*see* Methods) identified about 16,000 differentially methylated regions, and more than 80% of these DMRs were hypermethylated in the RYBP-deficient cells (Fig. 2a). YAF2 knockout showed no significant DMRs. The DNA hypermethylation after loss of RYBP often affected larger genomic regions that lost the PRC1 histone mark H2AK119ub1, as shown for example for the *HOXB* locus, a well-known Polycomb target (Fig. 2c,d). However, when considering CpG islands, the predominant targets of DNA hypermethylation in cancer, we observed that only 298 CGIs contained hypermethylated DMRs, which were abundant instead in intergenic and intragenic regions (Fig. 2b). Thus, RYBP inactivation alone was insufficient to cause extensive and specific CGI methylation as seen in tumors.

**Fig. 2.**
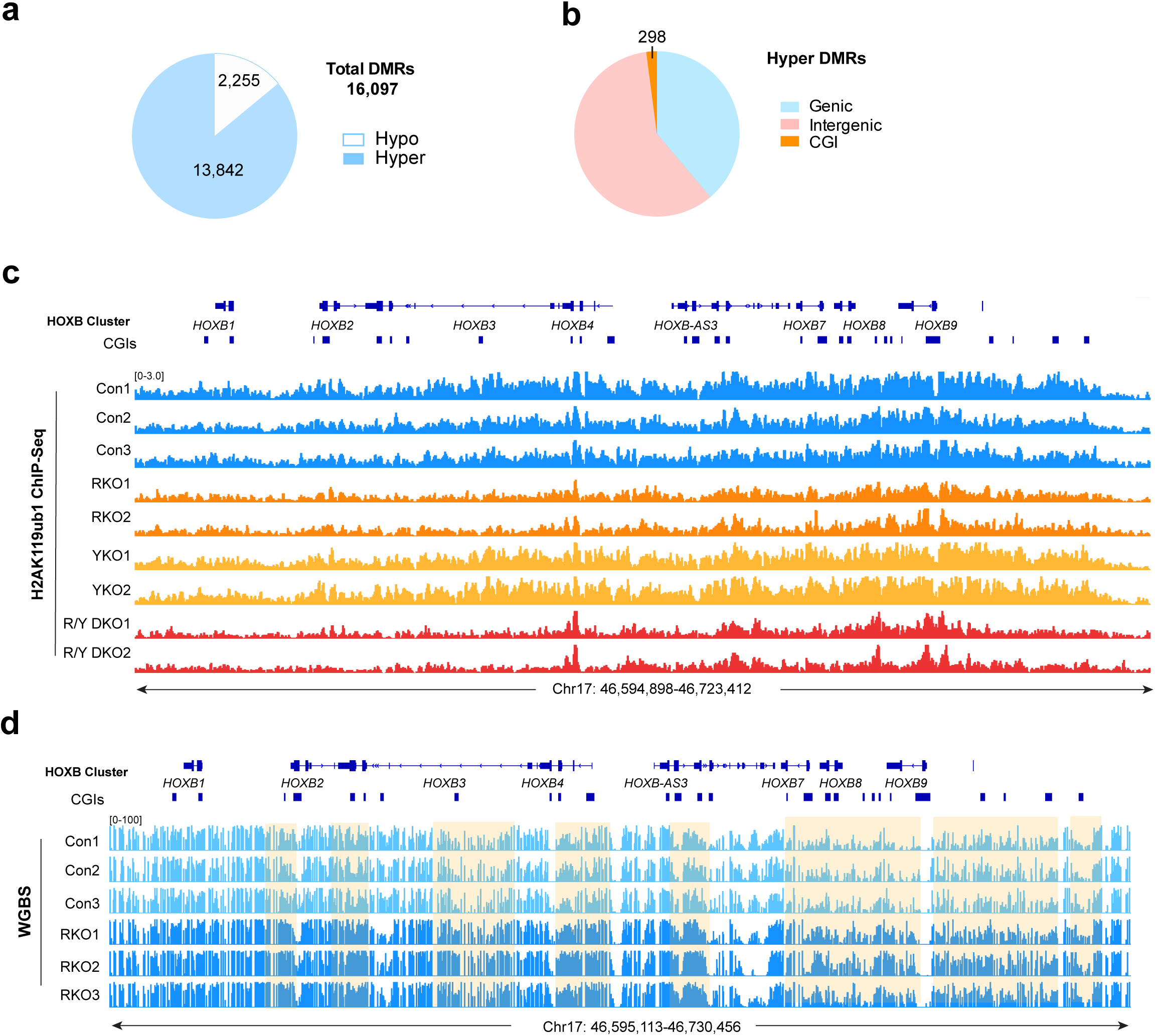
DNA hypermethylation in RYBP-deficient cells. **a.** Quantities of hypermethylated DMRs (Hyper DMRs) and hypomethylated DMRs (Hypo-DMRs) in RKO cells compared to controls. **b.** Quantities of hypermethylated DMRs that overlapped with the indicated genomic features in RKO cells compared to controls. **c.** IGV browser view of the *HOXB* cluster for H2AK119ub1 ChIP-seq data acquired from controls, RKO, YKO and R/Y DKO cells. **d.** IGV browser view of the *HOXB* cluster for WGBS DNA methylation data acquired from control and RKO cells. The hypermethylated regions are shaded.

## DNA hypermethylation after inactivation of RYBP and TET proteins

We next considered that 5mC oxidases, the TET proteins, can revert inappropriately incorporated 5mC back to the unmethylated state. TET1 and TET3 have been mapped to CpG islands, whereas TET2 has more commonly been associated with enhancer regions (Bogdanovic et al., 2016; Hahn et al., 2019; Hon et al., 2014; Jin et al., 2016; Wang et al., 2018; Williams et al., 2011). However, there is likely a considerable overlap between the three TET proteins in their functionality and genomic specificity. We first assessed the expression levels of the three *TET* genes in the parental bronchial cells and in the RYBP-inactivated cells using RNA-seq data. Expression of *TET2* and *TET3* was readily detectable but *TET1* levels in the control cells were extremely low (Extended Data Fig. 3a,b). After loss of RYBP, expression of *TET1* and *TET2* were significantly increased although *TET1* levels remained relatively low. Based on this data and our prediction that combined inactivation of RYBP and TET proteins could have a more pronounced phenotype, we proceeded to inactivate all three TET proteins in bronchial cells. We created single knockouts for TET2 and TET3, and various combinations of double and triple knockouts as illustrated by the clone tree (Extended Data Fig. 3d). The effort culminated in creation of a quadruple knockout that lacked RYBP, TET1, TET2, and TET3. For each targeting, three independent clones with biallelic frameshift mutations were created as confirmed by Western blotting for TET2 (Extended Data Fig. 3c), with no suitable antibodies available for human TET1 and TET3) and by extensive DNA sequencing (Supplementary Table 1; Extended Data Fig. 3d).

We performed WGBS on most of the derived knockout clones and determined DMRs (Supplementary Table 4, Excel file). A CIRCOS plot of DMRs for TET2 single KO, TET2/TET3 double KO (DKO), RYBP single KO, TET1/2/3 triple KO (TKO), and RYBP/TET1/2/3 quadruple KO (QKO) is shown in Figure 3a. Heat map analysis is shown in Figure 3b for total DMRs and in Figure 3c for CpG island DMRs. TET2 (and TET3) single knockout and TET2/TET3 DKO cells showed no DMRs that reached genome-wide statistical significance (Fig. 3a,d). DMRs in the RYBP single KO, the TET1/2/3 triple and the RYBP/TET1/2/3 quadruple knockouts were distributed uniformly along all chromosomes (Fig. 3a). Removing all three TET proteins led to more extensive DNA methylation changes and resulted in over 89,000 DMRs (Fig. 3d). The majority of DMRs in the TET TKO clones were hypomethylated, which is *a priori* unexpected for inactivation of 5mC oxidases. There is precedence however that TET protein loss leads to DNA hypomethylation, which was postulated to be caused by a redistribution of DNMT proteins upon loss of TET (Lopez-Moyado et al., 2019). Global DNA hypomethylation is observed in many cancer types and primarily involves heterochromatic regions. In the RYBP/TET1/2/3 QKO cells, we observed 71,000 DMRs, but now the direction of the DMRs was shifted again towards hypermethylation (Fig. 3a,d).

**Fig. 3.**
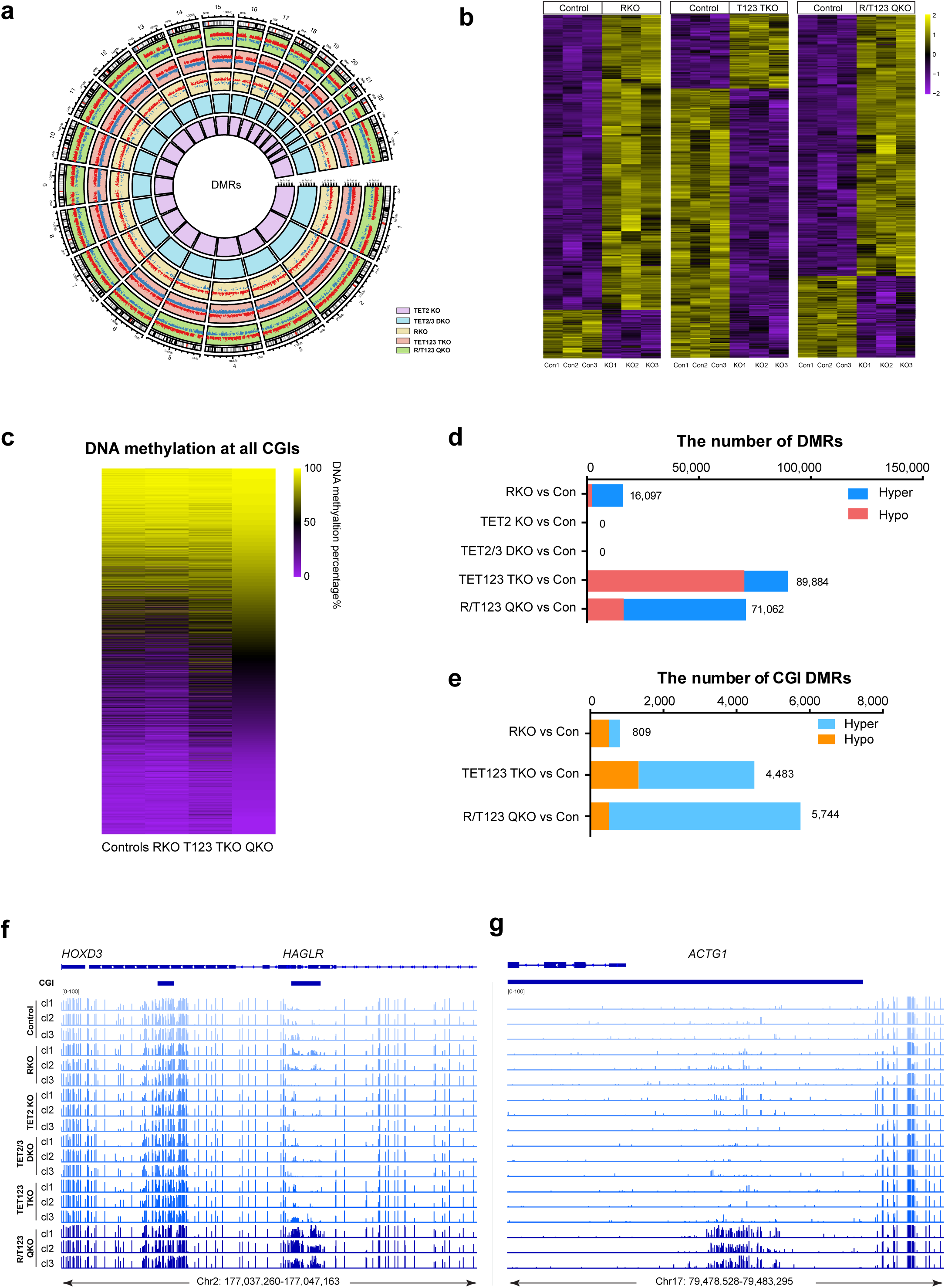
DNA methylation changes in cells deficient in RYBP and TET proteins. **a.** CIRCOS plot showing distribution of hypermethylation-DMRs and hypomethylation-DMRs on all chromosomes in RYBP KO, TET2 KO, TET2/3 DKO, TET123 TKO, and RYBP/TET123 QKO cells. Red dots denote hypermethylation-DMRs, blue dots denote hypomethylation-DMRs, and the coordinates represent the methylation percentage; positive values represent hypermethylation, negative values represent hypomethylation. Different background colors are used for the different knockout cells. **b.** Heatmap of DNA methylation changes (all DMRs in knockouts compared to controls) for RYBP KO, TET123 TKO, and RYBP/TET123 QKO. **c.** Heatmap showing DNA methylation at all CpG islands in control, RYBP KO, TET123 TKO and RYBP/T123 QKO. Each row represents one CpG island ranked by the methylation level in QKO. **d.** The numbers of all hypermethylated DMRs and hypomethylated DMRs in the RKO, TET2 KO, TET2/3 DKO, TET123 TKO, and R/TET123 QKO cells (knockouts compared to controls). **e.** The numbers of hypermethylated DMRs and hypomethylated DMRs that overlap with CGIs in the RKO, TET123 TKO, and R/TET123 QKO (knockouts compared to controls). The combined analysis of three independent clones is presented in panels a-e. **f, g.** IGV browser views of the *HAGLR* and *ACTG1* loci including CGIs for WGBS data acquired from controls, TET2 KO, TET2/3 DKO, TET123 TKO, and RYBP/TET123 QKO cells. Three independent clones in the different knockout cell lines are presented.

Focusing on CGIs, the TET triple knockouts and most significantly, the quadruple knockouts had predominantly hypermethylated regions, which affected over 5,000 CpG islands in the RYBP/TET1/2/3 QKO (Fig. 3e). This finding is different from the RYBP single knockout, which had more hypomethylated DMRs than hypermethylated DMRs at CGIs (Fig. 3e). The extent of DNA hypermethylation was most dramatic in the RYBP/TET123 QKO cells (Supplementary Table 5, Excel file). The RYBP/TET123 QKO had 3,176 hypermethylated CpG islands not found in either the RYBP single knockout or the TET1/2/3 TKO (Extended Data Fig. 4a). We show examples for the gene *HAGLR*, which is embedded in the *HOXD* locus (Fig. 3f) and for *ACTG1* (Fig. 3g). Making various pair-wise comparisons between cells of different genotype, we determined that many DMRs are common in these comparisons as shown in Extended Data Figures 4b and 4c. For example, comparing the DMRs found between QKO versus control, QKO versus TET triple KO, and QKO versus RYBP single knockout reveals that 1,018 CpG islands are in common for these comparisons.

## Hypermethylation of Polycomb targeted CpG islands

We intersected the Polycomb mapping data from our ChIP-seq experiments (Fig. 1; Supplementary Table 2) with the DNA methylation data obtained by WGBS. This analysis revealed that CGIs marked by H2AK119ub1 frequently become methylated in the RYBP/TET123 QKO cells (Fig. 4). Figure 4a shows several genome browser examples of methylation at CGIs associated with homeobox and other transcription factor genes. These CGIs are all marked by H2AK119ub1. We centered all CGIs of the genome in composite profile plots and observed an overall increase of DNA methylation levels in the QKO samples versus controls (Fig. 4b). We saw the same when we separately analyzed CGIs occupied by H2AK119ub1 (Fig. 4c), H3K27me3 (Fig. 4d) or RYBP (Fig. 4e). The increase of methylation affected the entire length of the CpG islands globally rather than being targeted to certain regions of them such as CGI centers or CGI shores (Fig. 4b-e).

**Fig. 4.**
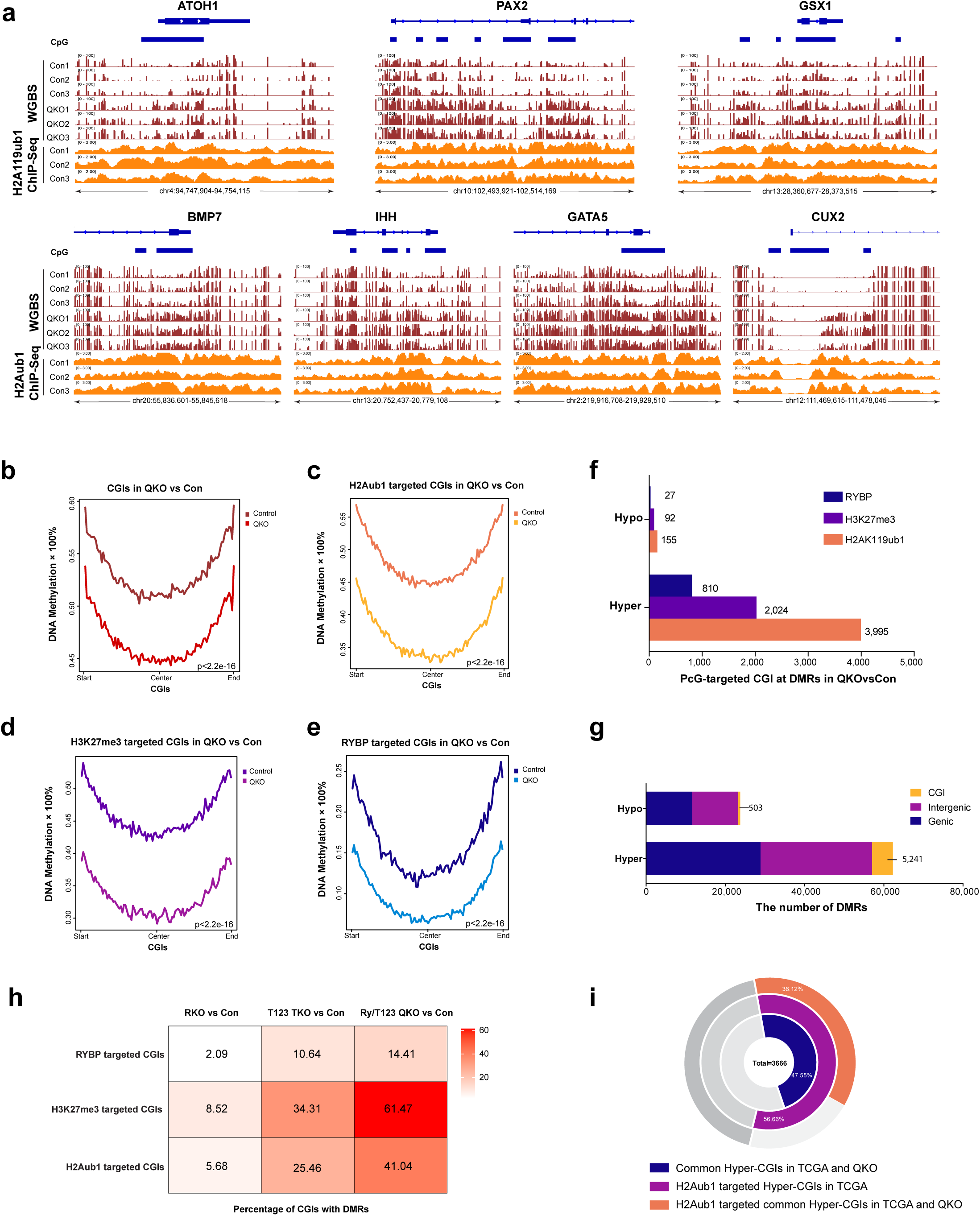
Hypermethylation of Polycomb target genes in the RYBP/TET1/2/3 knockout cells. **a.** IGV browser views of the genes, *ATOH1*, *PAX2*, *GSX1*, *BMP7*, *IHH*, *GATA5*, and *CUX2* showing WGBS data acquired from controls and R/T123 QKO cells and corresponding H2AK119ub1 ChIP- seq tracks. **b.** Metaplot illustrating DNA methylation levels at all CGIs in controls and R/T123 QKO cells. **c.** Metaplot illustrating DNA methylation levels at H2AK119ub1-targeted CGIs in controls and R/T123 QKO cells. **d.** Metaplot illustrating DNA methylation levels at H3K27me3-targeted CGIs in controls and R/T123 QKO cells. **e.** Metaplot illustrating DNA methylation levels at RYBP- targeted CGIs in controls and R/T123 QKO cells. *P* value is indicated in panels b-e. **f.** Numbers of RYBP-, H3K27me3-, and H2AK119ub1-targeted CGIs that contain Hyper-DMRs and Hypo-DMRs in R/T123 QKO cells relative to controls. **g.** Numbers of Hyper-DMRs that overlap with the indicated genomic features in R/T123 QKO compared to controls. **h.** Heatmap showing the percentage of all RYBP-, H3K27me3-, and H2AK119ub1-targeted CGIs that contain DMRs in RKO, TET123 TKO and R/T123 QKO cells, respectively. **i.** A total of 3666 hypermethylated CGIs from LUSC tumor samples are found in the TCGA dataset. The ring chart indicates the percentage of H2AK119ub1-targeted hypermethylated CGIs (Hyper-CGIs) in the TCGA dataset, and the percentage of them that overlap with Hyper-CGIs in the R/T123 QKO cells.

However, the numbers of CGIs that became hypermethylated were different depending on the associated mark (Fig. 4f). The largest number of CGIs subjected to hypermethylation in the QKO were those that carried the H2AK119ub1 mark (n=3,995), followed by H3K27me3 (n=2,024) and RYBP (n=810). Because the total number of CpG islands with hypermethylated DMRs in the QKO was around 5,200 (Fig. 4g), we calculated from these numbers that 76% of all CGI-specific hypermethylated DMRs were targeted to H2AK119ub1-marked CGIs. For comparison, in some cancer types, such as lung squamous cell carcinoma, 79% of the hypermethylated CGIs are associated with Polycomb target genes when H3K27me3 data from ES cells were used for comparison (Pfeifer and Rauch, 2009). We also determined the percentage of CGIs associated with a particular mark/protein that did undergo hypermethylation (Fig. 4h). This analysis showed that even though H3K27me3-marked targets are fewer in number (Fig. 4f), over 60% of them were methylated in the QKO. Forty-one percent of H2AK119ub1-marked CGIs became hypermethylated in the QKO (Fig. 4h). This number was smaller in the RYBP single KO (5.6%) and the TET-TKO (25%).

We wondered how the hypermethylation of CGIs seen in the QKO cells relates to DNA methylation changes seen in human lung cancers, in particular squamous cell carcinomas (SCC). We extracted the DNA methylation values at CGIs from the TCGA database and established a list of lung SCC hypermethylated CGIs for those samples for which normal lung controls were available (n=40). This list had 3,666 hypermethylated CGIs (Fig. 4i). Almost half of them (1,743 = 47.5%) overlapped with the hypermethylated CGIs found in the QKO (Fig. 4i; Supplementary Table 6, Excel file). A substantial fraction (56.6%) of the TCGA hypermethylated CGIs carried the H2AK119ub1 mark in bronchial cells. More than 1,300 CGIs (36.1%) carried the PRC1 mark and became hypermethylated in both the QKO and in human lung SCC (Fig. 4i). We acknowledge the inherent limitations of the TCGA data because “normal” lung controls represent a mix of many different cell types.

We wondered if the DNA hypermethylation events at CGIs in the RYBP/TET123 QKO and TET TKO cells were associated with specific DNA sequences. Motif analysis revealed the enrichment of transcription factor motifs in the hypermethylated regions (Extended Data Fig. 5). Most striking was the specific enrichment of homeobox transcription factor motifs, which occurred in both knockouts and were also seen when we subsampled the hypermethylated regions to focus on those marked by H2AK119ub1 (Extended Data Fig. 5a-c). These homeobox binding motifs bear same similarity, are generally AT-rich, and surprisingly lack any CpG dinucleotides (Extended Data Fig. 5a-c). However, there is a smaller set of CGIs that become hypermethylated but are *not* marked by H2AK119ub1 (about 1,000). Motif analysis of this subset showed the enrichment of TFs with motifs containing CG sequences, including, for example members of the ETS family of transcription factors, YY1, and E2F (Extended Data Fig. 5d). We do not currently know the reasons why such different motifs are found depending on Polycomb marking, but one possibility is that the non-CG motif-binding homeobox transcription factors are involved in recruiting PRC1 complexes at some early stage of development.

## Cell transformation and tumor growth of RYBP- and TET-deficient cells

The morphology of the TET-targeted cells resembled that of control cells but cells lacking RYBP had an enlarged cytoplasm (Extended Data Fig. 6). We performed cell growth assays in soft agar for the control cells, TET2 single KO, RYBP single KO, TET123 TKO and the RYBP/TET123 QKO cells (Fig. 5a-c). Confirming earlier data showing that HBEC3-KT cells are nontumorigenic (Ramirez et al., 2004; Vaz et al., 2017), the parental control cells did not grow in soft agar and neither did TET2 and RYBP single knockouts cells (Fig. 5a). TET123 TKO cells showed few, very small colonies. However, the RYBP/TET123 QKO cells showed high colony forming efficiency and generated large size colonies in soft agar, indicating that they had acquired properties of the transformed phenotype (Fig. 5a-c). This property was unrelated to an enhanced cell division rate because all TET- or RYBP-deficient cells grew more slowly than control cells (Fig. 5d).

**Fig. 5.**
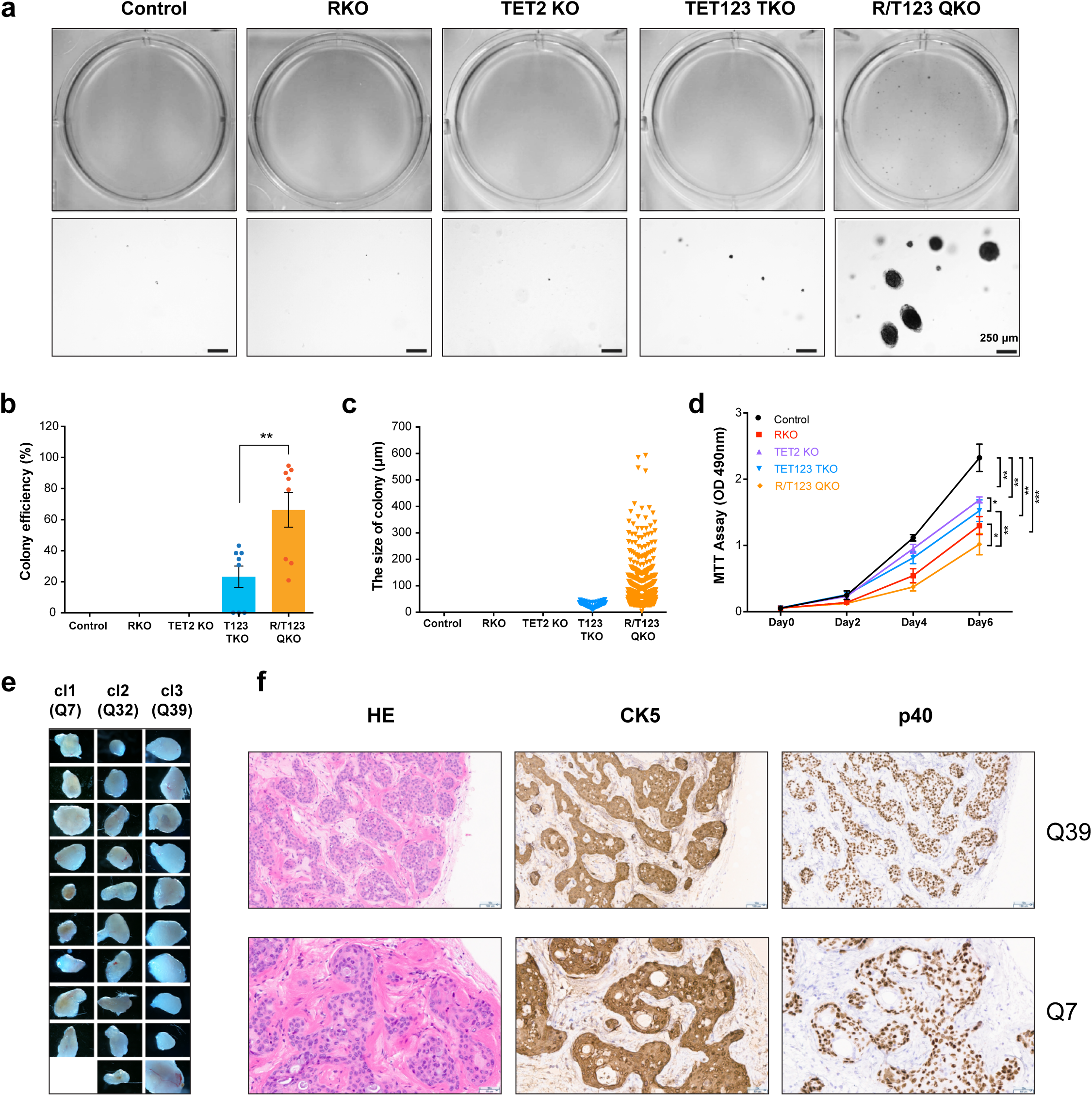
Growth properties of bronchial cells that lack RYBP and the TET proteins. **a.** Soft-agar assays to assess the colony forming potential of controls, RKO, TET2 KO, TET123 TKO, and R/TET123 QKO cells (top panel). Representative images of soft agar colonies are shown (bottom panel). Three replicates, three independent experiments, and three independent cell clones were used. Scale bars, 250 μm. **b** and **c.** Efficiency and quantities of colony formation as obtained in panel a. Date represents mean ± SE. \*\**p* < 0.01. **d.** Cell proliferation curves of controls, RKO, TET2 KO, TET123 TKO, and R/TET123 QKO cells. Three independent clones were tested in five replicates and two independent experiments. Date represents mean ± SD, \**p* < 10e^- 3^-10e^-10^, \*\**p* < 10e^-10^-10e^-15^, \*\*\**p* < 0.10e^-15^-10e^-25^. **e.** Images of xenografts from mice injected with R/T123 QKO cells. Ten or eleven injection sites were used for each cell clone. Magnification 1x. **f.** Representative histology of xenografts obtained from mice injected with R/T123 QKO cells (top row, clone Q39; bottom row, clone Q7). Parallel sections were stained from each xenograft. Representative sections stained with H&E, anti-CK5, and anti-p40 are shown. Scale bars, 100 µm, top row; 50 µm, bottom row.

To determine if growth in soft agar is related to tumor growth in mouse xenotransplantation experiments, we transplanted control and QKO cells subcutaneously into the flanks of immune-deficient female NSG mice. Two controls, wildtype cells and empty vector transformed wildtype cells, did not form tumors. Tumor formation was observed at 29 of 31 injection sites with the three QKO cell clones (QKO7, QKO32, and QKO39) (Fig. 5e). This result was repeated in an independent transplantation experiment. Histopathology review of QKO cell line xenografts showed varied amounts to tumor cells in nodular configuration and epithelioid clusters of tumor cells with small cystic structures, single cell necrosis, compressed and angulated cellular contours with invasive features and single cell migration suggestive of epithelial mesenchymal transition (EMT, most prominent for clone Q32). The cytologic features of tumors were most remarkable as the supporting matrix used in cell culturing (Matrigel-like material) was moderately well retained in the xenograft with minimal host (mouse) lymphoid or macrophage contaminant allowing for clean immunostaining. The clones QKO7 and QKO39 formed cell clusters of tumor cells with differentiation and cysts of variable size with single lining of tumor cells as determined by H&E staining (Fig. 5f), whereas QKO32 tumors presented with more isolated and solid clusters of cells and in streaming pattern at the xenograft periphery. (Extended Data Fig. 7d). We used immunohistochemical (IHC) methods to stain all QKO tumor samples with antibodies against the squamous cell carcinoma markers p40 (recognizing the short isoform of TP63 expressed from the *TP63* gene), and with antibodies against cytokeratin 5 (CK5). This staining showed high positivity suggesting that the tumors formed were squamous cell carcinomas (Fig. 5f). We also stained the sections with anti-vimentin (VIM) and SNAIL markers to determine if cells had undergone an epithelial to mesenchymal transition (EMT). The staining results suggested that clone Q32 has properties of EMT (Extended Data Fig. 7d). In Q32, although expression of *CDH1* (E-cadherin) was not downregulated by RNA-seq, several EMT marker genes were upregulated including *VIM*, *CDH2* (N-cadherin), *SNAI1/2*, *TWIST1/2 and TGFB2* and this was confirmed at the protein level and by immunofluorescence for VIM (Extended Data Fig. 7a-c).

## Activation of tumor-promoting signaling pathways

Since we observed efficient cell transformation for the R/TET123 QKO cells, we investigated if cancer-related pathways had become aberrant. We used RNA-seq to analyze gene expression patterns in the RYBP single knockout cells, TET123 TKOs and the QKOs (Supplementary Table 3, Excel file). The TET-TKO had the smallest number of differentially expressed genes (DEGs), only about 400 (Fig. 6a). Both the RYBP single knockout and the R/TET123 quadruple KO cells had over 2,000 DEGs. Many cancer-relevant genes were differentially expressed in the QKO versus control comparison. Some of these genes are listed in Figure 6a and genome browser views are shown in Figure 6b. There was a strong upregulation of genes of the AP-1 transcription factor family (*FOS*, *JUN*, *ATF3*, *FOSL1*, and others), upregulation of cell cycle genes such as cyclin D2 and *E2F1*, as well as downregulation of the cell cycle inhibitor, *CDKN2A* (Fig. 6b). KEGG pathway analysis (Fig. 6c) and GO analysis (Fig. 6d) showed the enrichment of cancer pathway-related terms, including Hippo signaling, Wnt signaling, TGF-beta signaling, and various other signal transduction pathways. Figure 6e shows the substantial upregulation of downstream growth-promoting genes negatively regulated by the Hippo pathway (*CCN2*, also called *CTGF*, *ANKRD1*, and *TGFB2*). To understand why these genes may be upregulated, perhaps in connection with the extensive DNA hypermethylation we had induced in the QKO cells, we looked for DNA methylation-associated gene silencing of potential regulators of these growth-enhancing pathways.

**Fig. 6.**
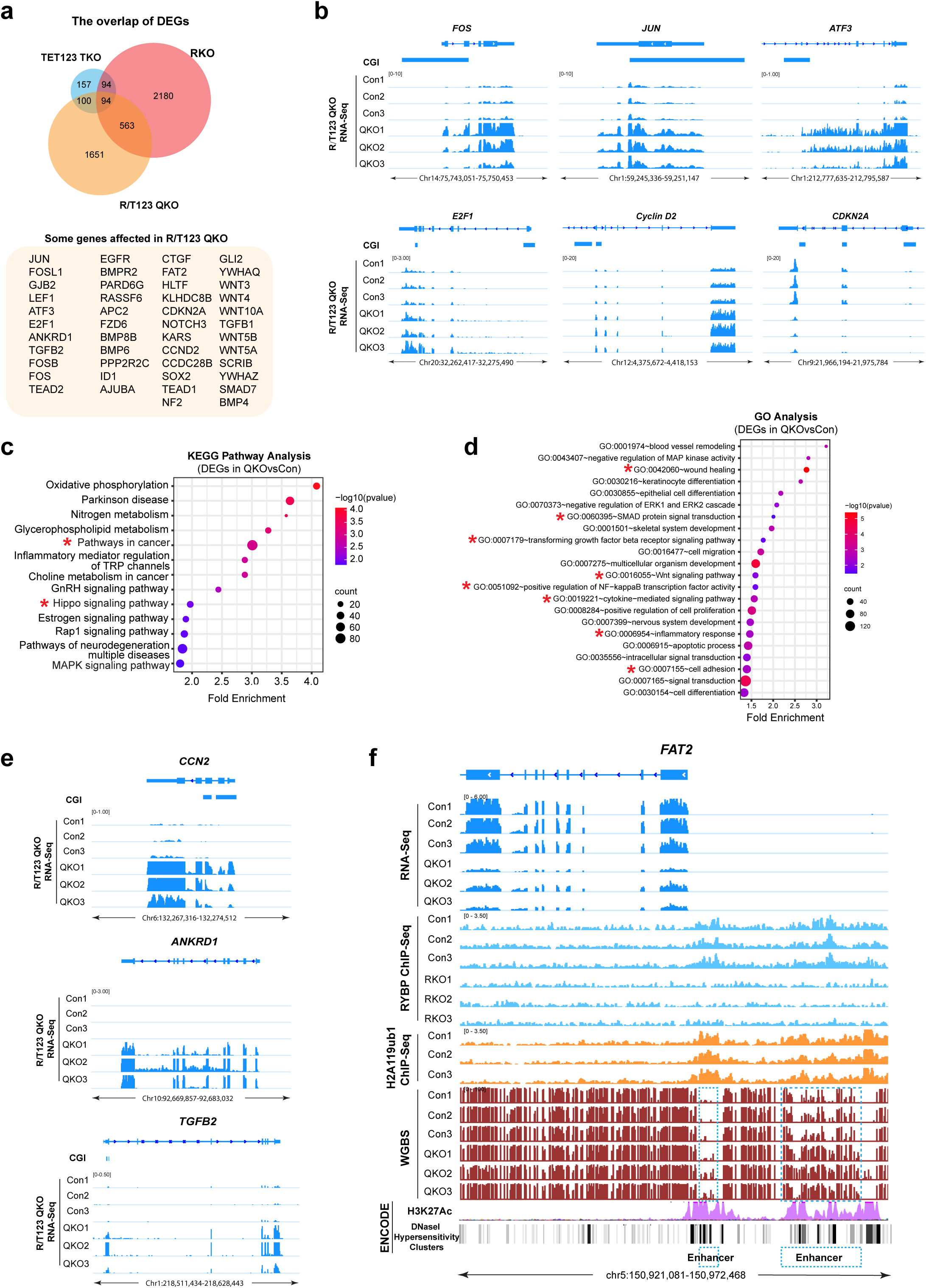
Alteration of cancer-related pathways in cells lacking RYBP and TET1/2/3. **a.** Overlap of differentially expressed genes in RKO, TET123 TKO, and R/T123 QKO cells. The combined analysis of three independent clones for each genotype is presented. Some examples of differentially expressed genes in the QKO vs. controls are listed. **b.** IGV browser views of the *FOS*, *JUN*, *ATF3*, *E2F1*, *Cyclin D2* and *CDKN2A* genes for RNA-seq data acquired from controls and R/T123 QKO cells. **c, d.** KEGG pathway and gene ontology (biological process) enrichment analysis of all differentially expressed genes in R/T123 QKO cells. **e.** IGV browser views of the *CCN2*, *ANKRD1*, and *TGFB2* loci for RNA-seq data acquired from controls and R/T123 QKO cells. **f.** RNA-seq, RYBP ChIP-seq, H2AK119ub1 ChIP-seq, and WGBS at the *FAT2* gene locus in controls, RKO or R/T123 QKO cells. H3K27ac ChIP-seq and DNasel hypersensitivity clusters from ENCODE are indicated at the bottom. The blue dashed lines indicate the promoter region and the upstream enhancer that become hypermethylated in the QKO.

First, we analyzed the correlation between methylation at promoter-associated CGIs and gene expression using global analysis (Extended Data Fig. 8). This analysis showed a relatively modest correlation. After applying a cutoff, we found 350 genes that were hypermethylated and downregulated (R=0.27, P=3.4e-07) (Extended Data Fig. 8a). For these downregulated genes, which in many cases carry both the H3K4me3 and H2AK119Ub1 marks in control bronchial cells, there is increased methylation at the TSS (Extended Data Fig. 8b). We show an example for the *GJB2* gene encoding a gap junction protein (Extended Data Fig. 8c), where methylation at the promoter-associated CGI coincides with strong downregulation of the gene.

Looking at known cancer-relevant genes, we did not find DNA hypermethylation of the CGI associated with the *CDKN2A* gene; instead, methylation was only seen in a region adjacent to that CGI. There are several negative regulators of the WNT pathway. However, we could not identify methylation-linked downregulation of any of them. For example, the Secreted Frizzled Related Protein genes (*SFRPs*) or *DKK* genes were either not expressed in bronchial cells (*DKK2*, *DKK4*, *DKKL1*, *SFRP2*-*5*) or not downregulated in the QKO (*DKK1*, *DKK3*, *SFRP1*).

We next analyzed upstream regulators of the Hippo pathway. FAT protocadherins are known positive regulators of Hippo anti-growth signaling (Gridnev and Misra, 2022). *FAT2* was strikingly downregulated in the QKO cells (Fig. 6f). According to ENCODE data, this gene is controlled by a proximal enhancer/promoter region and an upstream enhancer. Both regions were occupied by RYBP and by the histone mark H2AK119ub1. The two regions became hypermethylated in the QKO (Fig. 6f). In addition, we observed methylation and downregulation of several members of the *RASSF* gene family (*RASSF1A*, *RASSF4*, and *RASSF6*), which encode proteins that activate the mammalian MST1/2 Hippo kinases (Richter et al., 2009). The RASSF1A isoform is a known lung tumor suppressor (Dammann et al., 2000). We see methylation of its H2AK119ub1-marked promoter CpG island in the QKO and downregulation of the *RASSF1A* isoform (Extended Data Fig. 8d-f).

The RYBP/TET123 quadruple KO cells showed an upregulation of the *KRAS* gene (Extended Data Fig. 9a). There were no *KRAS* mutations in the QKO as determined by extensive DNA sequencing. The short arm of chromosome 12, where *KRAS* is located, showed a copy number gain in the QKO cells leading to upregulation of most genes on 12p (Extended Data Fig. 9b,c). We hypothesized that the 12p chromosomal instability in these cells may have been induced by the extensive DNA hypermethylation in these cells and by downregulation of genes that control genetic stability. Indeed, we found two genes that fulfill these criteria. The gene *HLTF* coding for helicase-like transcription factor, a RAD5-related translocase, is involved in genome stability and DNA repair (Poole and Cortez, 2017; van Toorn et al., 2022). This gene is downregulated in the QKO, its 5’ CpG islands is associated with RYBP and H2AK119ub1 in bronchial cells, and the CGI becomes hypermethylated in the QKO (Extended Data Fig. 9d). Another candidate gene for maintaining genome stability is *KLHDC8B*, which encodes a protein involved in centrosome function and chromosome stability (Krem et al., 2012). This gene is also strongly downregulated and becomes hypermethylated at the promoter CGI in the quadruple knockout cells (Extended Data Fig. 9e).

## DISCUSSION

We provide a mechanistic explanation for the origin one of the most pervasive epigenetic aberrations in human tumors, CpG island methylation, which occurs specifically at Polycomb target genes. CGI methylation in cancer is widespread, is seen in every malignancy, and is observed at a lower intensity already in premalignant tissues (Hanley et al., 2017; Hu et al., 2021; Tommasi et al., 2009; Wijetunga et al., 2016; Wijetunga et al., 2017) and even in tissues undergoing inflammatory reactions and during aging (Hahn et al., 2008; Maegawa et al., 2010). We propose that Polycomb-marked CGIs are inherently susceptible to hypermethylation owing to long-term instability of protection mechanisms, as previously proposed for aging tissues (Jung and Pfeifer, 2015). The methylation increase is dramatically accelerated during malignant progression, eventually resulting in several thousand of such CGIs becoming methylated in tumors.

Various mechanisms for the CGI hypermethylation process have been proposed. They include, for example overexpression of DNMT proteins in cancer, but this hypothesis has not been fully substantiated. DNMT targeting might be involved, for example via the recently described interaction of the unstructured N-terminus of DNMT3A1 and H2AK119ub1 (Gu et al., 2022). Other studies have shown alterations in chromatin configuration, such as the erosion of H3K4me1 at CpG island borders, that can lead to DNA hypermethylation (Skvortsova et al., 2019). We focused on protection mechanisms that may break down over time and during malignant transformation. These protective shields against methylation are two-pronged, consisting of the Polycomb complexes themselves and of the 5mC oxidases. Here we examined the role of the RYBP protein because it is a critical activator of variant PRC1 complexes, and unlike most other Polycomb genes, *RYBP* is downregulated in many tumor types (Fig. 1a, Extended Data Fig. 1a). However, this choice does not mean that RYBP dysfunction is the only possible defect of the Polycomb system in cancer. Other members of PRC1, or even PRC2, may also become dysfunctional through various mechanisms, such as downregulation of expression (e.g., *PCGF5*, *CBX7*), increased protein instability, or altered posttranslational modifications. Furthermore, simultaneous dysfunction of several members of these large complexes may be additive.

Cells contain a powerful backup system to correct inappropriately introduced 5- methylcytosines by demethylation mediated by the TET proteins. TET1 and TET3 are known to bind to CGIs, likely through their CXXC domains, which have affinity to unmethylated CpG-rich DNA sequences (Jin et al., 2016; Williams et al., 2011). TET1 has been found in association with Polycomb complexes and binds to Polycomb target genes (Williams et al., 2011; Wu et al., 2011). Also, TET2, which is often mapped to enhancers, may be recruited to CGIs by its associated binding partner CXXC4/IDAX, which is another CXXC domain family protein (Ko et al., 2013). This considerable redundancy helps to protect CpG islands from methylation. However, TET protein function appears to be substantially reduced in tumors. First, TET2 is mutated in hematological malignancies. Mutations in isocitrate dehydrogenase (IDH1, IDH2) create a neomorphic enzyme that generates 2-hydroxyglutarate, a competitive inhibitor of TET proteins (Dang et al., 2009). However, IDH mutations are limited to certain tumor types such as gliomas and cholangiocarcinomas (Pirozzi and Yan, 2021). On a much broader scale, TET protein activity appears to be strongly diminished in all human solid tumors, as shown by vanishing levels of 5- hydroxymethylcytosine in the malignant tissue, even in nondividing cells of the tumor mass (Jin et al., 2011). It has been puzzling to understand the cause of this 5hmC loss, but this finding is not simply related to downregulation of *TET* genes in tumors. TET protein function may be impaired by other cancer-associated disturbances such as changes in metabolism that affect the availability of the co-factors alpha-ketoglutarate and/or ascorbic acid, and/or a hypoxic tumor environment (Thienpont et al., 2016). Regardless of the exact mechanisms for loss of TET function in cancer, our data shows that inactivation of TET proteins in lung epithelial cells in combination with a dysfunctional PRC1 complex leads to extensive DNA hypermethylation of Polycomb target genes, mimicking the situation commonly observed in most human cancers.

The second major finding of this work is identifying the consequences of DNA hypermethylation of Polycomb target genes in tumors. We detected the conversion of nontumorigenic bronchial cells to transformed cells that show anchorage-independent growth and form tumors in immune-deficient mice without the introduction of cancer-driving mutations. In previous studies, these HBEC3-KT cells could not even be transformed by simultaneous introduction of a *KRAS* mutation, *EGFR* mutation and by inhibiting the TP53 tumor suppressor (Sato et al., 2006). *TET* genes are mutated in a small percentage (<10%) of human lung tumors. A recent study observed tumor formation in a mouse model with *Kras* mutation and loss of *Tet* genes, but the induced hypermethylation events were not targeted to CpG islands (Xu et al., 2022). Although we observed a 1.9-fold increase of *KRAS* expression due to a copy number gain of chr12p, there were no *RAS* gene mutations, and we favor the model that the disturbance of several other cancer-relevant signaling pathways was due to direct epigenetic perturbation. We observed a reduced output of the Hippo tumor suppressor pathway as shown by strong upregulation of canonical Hippo target genes such as connective tissue growth factor (*CTGF*) (Fig. 6). Another characteristic finding in our study was the increased expression of AP-1 genes, which may be a critical event in certain types of lung cancer (Kim and Minna, 2022). The TEAD transcription factors that are negatively controlled by the Hippo pathway often co-localize and collaborate with AP-1 proteins, and this combined activity drives cell proliferation (Tang et al., 2022; Zanconato et al., 2015). We found several induced CGI hypermethylation events that have the potential to be cancer drivers. These methylation target genes were positive regulators of the Hippo anti-growth pathway (*RASSF* genes and *FAT2*) that became downregulated in association with promoter methylation and several genes controlling genomic stability. The *FAT2* gene, encoding an atypical protocadherin and positive regulator of the Hippo pathway (Fulford and McNeill, 2020), is also mutated in close to 10% of non-small cell lung cancers (COSMIC database) and this gene is highly expressed in human basal respiratory cells of the bronchi (proteinatlas.org). In lung cancers, expression of *FAT2* is very heterogenous between patients with many having close to zero expression. It would be difficult to dissect which one of these hypermethylation events is the most consequential one for tumor formation because they co-occur upon misregulation of the two epigenetic pathways. We propose that a combination of these DNA hypermethylation events and the ensuing changes in gene expression, brought about by our specific manipulations of the epigenome, effectively overcome an anti-proliferative barrier eventually leading to cell transformation.

In summary, based on our findings in non-transformed lung epithelial cells, a clear model emerges for the formation of lung cancer, as presented in Extended Data Figure 10. Further studies will test whether these mechanisms can explain the initiation of other human cancers.

## Materials and Methods

### Cell Culture

Human bronchial epithelial cells (HBEC-3KT, ATCC, CRL-4051) were cultured in keratinocyte serum-free medium (KSFM; Thermo, 17005042) containing 50 mg/ml of bovine pituitary extract (BPE) and 5 ng/ml of human recombinant epidermal growth factor (rEGF). Cells were incubated at 37 °C with 5% CO_2_. The media was changed every 3 to 4 days.

### Generation of HBEC3 knockout cell lines using CRISPR/Cas9

CRISPR gRNAs were designed to target the genomic sequence, and to introduce frameshift mutations into exon 2 of RYBP and YAF2, or the N-terminal region of the catalytic domains for TET1, TET2 and TET3. All CRISPR gRNAs used in this study are listed in Supplementary Table 7.

### Lentivirus production

The pLentiCRISPR E plasmid (Addgene, 78852, a gift from Phillip Abbosh) was modified with different selectable marker genes, blasticidin, hygromycin or with EGFP, then co-transfected into HEK293T cells with lentiviral packaging plasmids psPAX2 and pMD2.G (Addgene, 12260 and 12259, gifts from Didier Trono). HEK293T cells were cultured in DMEM plus 10% fetal bovine serum and seeded in 6-well cell plates the day before transfection in such way that they would be 80%-90% confluent at the time of transfection. For transfection of one well, 20 µl of FuGene HD Transfection Reagent (Promega, E2311) was diluted into 500 μl of Opti-MEM and then the following DNAs were added: 1 μg gRNA vector, 0.75 μg psPAX2, and 0.25 μg pMD2.G. The solution was briefly vortexed and incubated at room temperature for 20 minutes. The mixture was then gently added to 6-well plates with 1.5 ml DMEM. All media was aspirated after 10 hours and replaced with fresh DMEM containing 10% FBS. Viral particles were harvested 48 hours after this media change and frozen at -80°C.

### Identification of mutant clones

Two days after the virus infection, blasticidin (4 μg/ml, ThermoFisher, A1113903) or hygromycin B (40 μg/ml, ThermoFisher, 10687010) were added into the medium for selection for one week. For EGFP selection, the cells were cultured for two weeks until cells with visible green color appeared. For cells with no selection marker, the cells were cultured for two weeks. HBEC cells were dissociated into single cells and replated at 500-1,000 cells per 10 cm dish. Cells were allowed to grow until colonies from single cells became visible (10-14 days), then the single colonies were manually picked under a microscope and distributed into 24-well plates. The cells were kept under antibiotics. Colonies were expanded, and 30 to 40 plasmid sequences for each colony were analyzed by Sanger sequencing at gRNA-targeted positions. Sequencing primers are shown in Supplementary Table 7. For each genotype of HBECs, we derived three independent cell clones. The genotype of each clone is listed in Supplementary Table 1 and in Extended Data Figure 3. The evolutionary tree of all cell clones is shown in Extended Data Figure 3.

### RNA isolation, mRNA-seq and data analysis

Total RNA was isolated with PureLink RNA Mini Kit (Invitrogen, 12183020) from HBEC controls and various HBEC targeted clones (three different clones for each genotype). RNA quality was assessed by the Agilent Bioanalyzer 2100, and RIN scores of all samples were above 9.90. We performed RT-PCR as described (Dammann et al., 2000). Briefly, 2 μg of RNA for each sample was converted to cDNA with gene-specific primers using SuperScript III Reverse Transcriptase Kit (Invitrogen, 18088). mRNA level was determined with pfuUltra II Fusion HS DNA polymerase (Agilent, 600670). PCR primers are listed in Supplementary Table 7. For mRNA-seq, the libraries were prepared from total RNA with the KAPA RNA HyperPrep kit with RiboErase (KAPA Biosystems). Library size distributions were validated on the Bioanalyzer (Agilent Technologies), then the libraries were sequenced using an Illumina NextSeq500 system with 75 bp single read runs at the Van Andel Institute Genomics Core.

For mRNA-seq data analysis, 75-bp single-end reads were trimmed with Trim Galore (version 0.4.0). Reads were aligned to the human genome hg19 with STAR (version 2.5.1), then gene count was performed with STAR. Differential gene expression was determined with the Limma (version 3.38.2) statistical package as descried previously (Huang et al., 2021). Differential expression *P* values were adjusted for multiple testing correction using the Benjamini-Hochberg method in the stats package. . Statistical significance for differentially expressed genes was fold change > 2 with *q* < 0.05. Heatmaps were generated with pheatmap (version 1.0.12).

### ChIP-seq and data analysis

For RYBP ChIP (antibody from Millipore, AB3637), cells were fixed with 2.5 mM EGS (ethylene glycol-bis-succinimidyl-succinate) (Thermo Scientific, 21565) for 45 minutes, then washed twice with PBS and fixed again with 1% formaldehyde in 50 mM HEPES-KOH, pH 7.5, 100 mM NaCl, 1 M EDTA, pH 8.0, 0.5 mM EGTA, pH 8.0, for 8 minutes. For H2AK119ub1 ChIP (Cell Signaling Technology, antibody 8240) and H3K27me3 ChIP (Cell Signaling Technology, antibody 9733), cells were crosslinked with 1% formaldehyde only, for 8 minutes, then fixation was stopped by adding glycine at 2.5 M concentration for 5 minutes. Fixed cells were suspended in lysis buffer (50 mM HEPES-KOH, pH 7.9, 140 mM NaCl, 1 mM EDTA, 10% glycerol, 0.5% NP40, 0.25% Triton X-100, protease inhibitors) for 10 minutes on ice. Then the cells were pelleted and re-suspended with wash buffer (10 mM Tris-Cl, pH 8.1, 200 mM NaCl, 1 mM EDTA, pH 8.0, 0.5 mM EGTA, pH 8.0, protease inhibitors) and shearing buffer (0.1% SDS, 1 mM EDTA, 10 mM Tris-Cl, pH 8.1, protease inhibitors). The cell pellet was resuspended with 1 ml of shearing buffer, and the DNA was sheared to 300 to 500 bp size fragments with a Covaris E220 Evo sonicator. Twenty micrograms of chromatin, antibody and protein-G bead complexes were incubated overnight on a rotator at 4°C. Then, the beads were washed with low salt buffer (0.1% SDS, 1% Triton X-100, 2 mM EDTA, 20 mM HEPES-KOH, pH 7.9, 150 mM NaCl), high salt buffer (0.1% SDS, 1% Triton X-100, 2 mM EDTA, 20 mM HEPES-KOH, pH7.9, 500 mM NaCl) and LiCl buffer (100 mM Tris-Cl, pH 7.5, 0.5 M LiCl, 1% NP40, 1% sodium deoxycholate) twice and with TE buffer (10 mM Tris-Cl, pH 8.1, 1 mM EDTA) once. Purified DNA was quantified for library preparation.

TruSeq ChIP Sample Preparation Kit (Illumina, IP-202-1012, IP-202-1024) was used according to the manufacturer’s instructions to perform ChIP-seq library preparation. Briefly, 5 ng of DNA was used as starting material for input and IP samples. Libraries were amplified using 14 cycles on the thermocycler. Libraries were validated using the Agilent High Sensitivity DNA Kit. Then libraries were submitted to the Van Andel Institute Genomics Core for sequencing with an Illumina NextSeq500 sequencer with 75 bp single read runs.

75-bp single-end reads were trimmed with Trim Galore (version 0.4.0) with default parameters, then the trimmed reads were mapped to human genome hg19 with Bowtie2 (version 2.3.5). After deduplication with the software picard (version 2.19.0), the deduplicated reads were used for peak calling by HOMER (version 4.10) (Heinz et al., 2010) with the following settings: -B -region -size 1000 -minDist 2500 -P 0.01 -F 2.0 -L 2.0 -LP 0.01. Peak motifs were identified with HOMER based on the peak calling information. Output bedgraph files from HOMER were processed into bigwig files with UCSC Exe Utilities bedGraphToBigWig (version 1.04.00) (Kent et al., 2010), then bigwig files were loaded into IGV for genome browser snapshots of ChIP-seq enrichment (Thorvaldsdottir et al., 2013). Density plots of ChIP-seq enrichment at peak center regions were made with R package genomation (version 1.28.0) (Akalin et al., 2015). Heatmap for comparing the enrichment pattern between control and knockout samples were plotted with R package Enrichedheatmap (version 1.22.0) (Gu et al., 2018), which are presented in Figure 1C. Peaks were sorted by the peaks in control. For all overlap analysis of ChIP peaks, we used bedtools - intersection function, as well as for all other region overlap analysis in this study. Significant difference between two groups in metaplot was calculated by Welch Two Sample t-test with R (version 4.2.0).

### WGBS and data analysis

Genomic DNA was extracted from HBEC controls and various HBEC derivative cells (three different clones for each group) using a Quick-DNA Miniprep Plus kit (Zymo Research, D4070). DNA samples were submitted to the Van Andel Institute’s Genomics Core (Controls, RYBP KO, TET2 KO, TET2/3 DKO, TET1/2/3 TKO, and RYBP/T123 QKO) or Fulgent (YAF2 KO and TET3 KO) for library preparation. The Accel-NGS Methyl-Seq DNA Library Kit (Swift Biosciences, Ann Arbor, MI, 30024) was used according to the manufacturer’s instructions to perform bisulfite conversion and library preparation. Sequencing was performed with an Illumina NovaSeq 6000 or HiSeq 2500 system with 150 bp paired end read runs. Generally, all libraries displayed high Q-scores (>30) in both read pairs. We obtained data corresponding to approximately 24x genome coverage on average. Paired-end sequencing reads were aligned to the hg19 human genome using Bismark (Krueger and Andrews, 2011). Adaptors and low-quality reads were trimmed using the parameters as described previously (Huang et al., 2021).

We used DMRseq (version 0.99.0) (Korthauer et al., 2019) to identify differentially methylated regions (DMRs). CpG methylation values were called by the Bismark methylation extractor script provided with Bismark, using the parameters as described previously (Huang et al., 2021). Sequencing depth for CpGs with at least three covering reads for each sample were considered for DMR calling, as well as for further analysis of DNA methylation. A single CpG coefficient cutoff of 0.05 was used for candidate regions. Significant DMRs between the control and modified cells were identified using q < 0.05.

To analyze the DMR motifs, HOMER was utilized for identifying putative motifs within DMRs. We used the following parameters on bed files of DMRs: findMotifsGenome.pl -size given -mask -cpg. Known motifs (as opposed to de novo motifs) from HOMER are presented in Extended Data Figure 5. We identified the functional enrichments of DMRs and plotted them with ReactomePA (version 1.40.0) (Yu and He, 2016), ChIPseeker (version 1.32.0) (Yu et al., 2015) and the clusterProfile (version 4.4.4) package (Wu et al., 2021). To identify the DMRs among different genotype groups, upset plots and pairwise plots are presented in Extended Data Figure 4 as plotted by Python tool Intervene (Khan and Mathelier, 2017), with intervene upset and intervene pairwise function. IGV was used to view the data tracks. We plotted the heatmaps of DMRs and CGIs among different genotype groups with R package pheatmap (version 1.0.12).

The CpG densities of methylation distribution between samples were calculated and plotted by the R package genomation (version 1.28.0). We plotted the Circos plot of DMRs of different genotype groups with circlize (version 0.4.10) (Gu et al., 2014).

Significant difference between two groups in metaplots was calculated by Welch Two Sample t-test with R (version 4.2.0).

The sequencing copy number on Chromosome 12 was visualized with the GenVisR (version 1.28.0.) (Skidmore et al., 2016).

### Gene ontology and pathway enrichment analysis

We used various gene sets for gene ontology and pathway enrichment analysis of differential expressed genes using DAVID functional annotation analysis (Huang da et al., 2009). Functional enrichment of ChIP-peaks was performed with R package ReactomePA (version 1.40.0) (Yu and He, 2016), ChIPseeker (version 1.32.0) (Yu et al., 2015), and clusterProfile (version 4.4.4) package (Wu et al., 2021).

### Nuclear protein and histone extraction and Western blotting

We isolated nuclear proteins with NE-PER™ Nuclear and Cytoplasmic Extraction Reagents kit (Thermo, 78835) according to the manufacturer’s instructions.

Histone acid extraction was performed as previously reported (Shechter et al., 2007). Briefly, the cell pellet from one 10 cm dish was washed with PBS twice, then resuspended and incubated with 1 ml hypotonic lysis buffer including additionally 50 μM of the deubiquitinase inhibitor PR-619 for 30 minutes on a rotator at 4°C. We resuspended the nuclei in 0.4 N H_2_SO_4_ and incubated on a rotator overnight. We added trichloroacetic acid to the supernatant containing histones and incubated the solution on ice for 30 minutes. We collected precipitates by centrifugation. Before dissolving the histone pellet in water, we performed two steps of washing with ice-cold acetone.

The antibodies for Western blotting are listed in Supplementary Table 8. We used Image J for quantitative analysis of Western blot signals of H2AK119ub1, H3K27me3 and histone H3.

### Immunofluorescence and microscopy

Cells were seeded in a chamber and fixed with ice-cold 100% methanol for 15 minutes at -20°C, then rinsed three times in PBS for 5 minutes each. Then, we blocked the samples with 5% BSA in PBST (PBS containing 0.3% Triton X-100) for 1 hour, and subsequently incubated them with the diluted primary antibodies in 1% BSA in PBST for overnight at 4°C. Cells were washed three times with PBS, incubated with fluorescent secondary antibody in 1 % BSA for 1 hour at room temperature in the dark, and finally decanted and washed in the dark prior to taking images. We captured fluorescent images on a Leica inverted microscope.

### Cell proliferation and soft agar growth assays

We monitored cell proliferation using MTT assay (Sigma, M5655). One thousand viable cells were seeded into 96-well plates, and cells were incubated with MTT solution in a CO_2_ incubator for 2 hours. We added 150 microliters of DMSO to each well until all crystals were dissolved. We measured the absorbance at 570 nm on a plate reader (Biotech Instruments, USA).

We determined the clonogenic capacity of HBEC3 and modified cells by soft agar assays. We suspended 5,000 viable cells and seeded them in 0.35% agar in K-SFM medium in 6-well plates and overlaid them with a 0.5% agar base in the same medium. We stained the colonies with 0.02% iodonitrotetrazolium chloride (0.02% INT, Sigma, I10406) and counted them using a microscope.

### In vivo tumorigenicity and histology analysis

We used female NSG mice (4-5 weeks old) (Jackson Laboratories) for these studies. Mice were housed at Van Andel Institute animal care facility. All animal experiments were approved by the Van Andel Institute Animal Care and Use Committee. All animal care and use protocols followed were in accordance with guidelines of the Institutional Animal Care and Use Committee (IACUC). We injected mice subcutaneously in the left and right flanks with 6x 10^6^ viable control cells or QKO cells in 0.2 ml of KSFM with Cultrex BME (R&D Systems, 3632-010-02). Mice were monitored every week for tumor formation for up to two months. Formalin-fixed, paraffin-embedded (FFPE) xenograft tumor tissue was sectioned and stained with hematoxylin and eosin (H&E) for histologic analysis. We performed immunohistochemical staining for CK5, p40, SNAIL and VIM at the Van Andel Institute Pathology and Biorepository Core with antibody specifics in Supplementary Table 8. Immunohistochemical staining was performed on an automated immunostainer (Dako) using standard protocols and stained slides were scanned to digital files (Aperio Scanscope).

### TCGA data analysis

For analysis of Infinium 450K methylation data from TCGA, we downloaded the raw IDAT files by R Bioconductor package TCGAbiolinks (version 2.24.3) (Colaprico et al., 2016). We used R Bioconductor package minfi (version 1.42.0) (Aryee et al., 2014) to process the raw IDAT files. The function preprocessNoob was applied for background correction based on the normal-exponential out-of-band (NOOB) background subtraction method (Triche et al., 2013), and then preprocessFunnorm function was used for further processing by functional normalization method (FunNorm) (Fortin et al., 2014). We used the object containing Beta-values returned by the preprocessFunnorm function for DMR calling by bumphunter (version 1.38.0) (Aryee et al., 2014) with bump hunting algorithm (Jaffe et al., 2012)s, with the cutoff 0.1 for the Beta-values and a large number of permutations, B=1000. For the downstream analysis, only the DMRs with at least two probes in a minimal region of 300 bp were considered. For RNA expression analysis of TCGA data, we downloaded the expression data by FirebrowseR (version 1.1.35), then generated box plots for comparing normal solid tissue and primary tumor with R (version 4.2.0).

### Statistical analysis

Data are presented as mean ± SD (unless otherwise noted) and were derived from at least three independent experiments. Data on replicates is given in the Figure Legends. Statistical analysis was performed using the two-tailed t-test or one-way comparison ANOVA. Statistical analyses for DNA methyl-seq, RNA-seq, and ChIP-seq data are described in those respective methods sections.

### Data availability

All sequencing datasets are available under the Gene Expression Omnibus accession number GSE208689.

## Supplemental information

Supplemental information includes Tables for oligonucleotides and antibodies and Excel files summarizing data analysis and can be found with this article.

## Acknowledgements

We thank Marie Adams, Marc Wegener, Rebecca Siwicki, and Katelyn Becker from the Van Andel Institute’s Genomics Core, Sidney Hein and Lisa Turner from the Pathology and Biorepository Core, the Microscopy Core, and Adam Rapp and personnel from the Vivarium for their expertise and support. We are grateful to Piroska Szabó for critical reading of the manuscript.

This work was supported by NIH grant CA234595 to GPP.

## Author contributions

W.C. carried out most of the cell culture, cloning, in vitro and in vivo experiments, methylation assays, gene expression, and ChIP sequencing studies with advice and assistance from S.-G.J. and J.J. Z.H. carried out most of the bioinformatics analyses, with assistance from W.C. G.H. performed analysis of tumor and histology sections. G.P.P conceived the study and designed experiments, supervised the work, and participated in data analysis. W.C. and G.P.P. wrote the manuscript with assistance from Z.H. All authors participated in editing the manuscript.

## Conflicts of interest

The authors declare no conflicts of interest.

## SUPPLEMENTARY TABLES

**Supplementary Table 1.** List of HBEC3 clonal KO cell lines. Mutated DNA sequences are indicated.

**Supplementary Table 2. ChIP-seq peaks.**

Excel file.

List of the RYBP, H2AK119ub1 and H3K27me3 peaks in controls determined by ChIP-seq. Hg19 genomic coordinates are indicated. The combined analysis of three independent clones in controls and two independent clones in knockout clones is presented.

**Supplementary Table 3. Differentially expressed genes.**

Excel file.

Differential gene expression analysis results for RKO, R/Y DKO, TET1/2/3 TKO, and RYBP/TET123 QKO compared to control HBEC3 cells. RKO, R/Y DKO and controls (5 different clones) from one run of mRNA-seq are presented. RKO, TET123 TKO, R/T123 QKO and controls (3 different clones) from another run of mRNA-seq are presented. Genes are sorted by fold change (1.5 or 2) of expression (KOs versus controls). Ensembl, gene symbol, log-transformed fold change, *P* value and adjusted *P* value (Benjamini-Hochberg correction) are provided. The combined analysis of three independent clones in different knockout cell lines is presented.

**Supplementary Table 4. DMRs.**

Excel file.

DMRs for RKO, TET1/2/3 TKO, and RYBP/TET123 QKO compared to controls derived from WGBS analysis. DMRs of different genotypes are separated into different sheets for hypermethylated

DMRs and hypomethylated DMRs. Hg19 genomic coordinates and CpG islands are indicated. The combined analysis of three independent clones is presented.

**Supplementary Table 5. Hypermethylated DMRs in the RYBP/TET1/2/3 quadruple knockout and their associated genes.**

Excel file.

Hypermethylated DMR regions of RYBP/TET123 QKO that intersect with CGIs and associated genes are listed. Hg19 genomic and gene symbol coordinates are indicated.

**Supplementary Table 6. List of the CGIs that overlap between the TCGA dataset of lung squamous cell carcinoma (LUSC) and RYBP/T123 QKO cells.**

Excel file.

List of the CGIs that overlap between the TCGA dataset of lung squamous cell carcinoma (LUSC) and RYBP/T123 QKO cells. Hg19 genomic coordinates are indicated. Almost half of them (1,743 = 47.5%, from TCGA) overlapped with the hypermethylated CGIs found in the QKO.

**Supplementary Table 7. List of DNA oligonucleotides.**

List of gRNA Target Sequences. Guide RNAs were used for targeting of the *RYBP*, *YAF2*, *TET1*, *TET2* and *TET3* genes. Listed are the oligonucleotides used for genotyping to identify knockouts of *RYBP*, *YAF2*, *TET1*, *TET2* and *TET3*, and PCR primers for amplifying *RASSF1A*.

**Supplementary Table 8. List of antibodies.**

Antibody target, suppliers, catalogue number, assay, and amount or dilution used are indicated.

---

**Extended Data Fig. 1.**
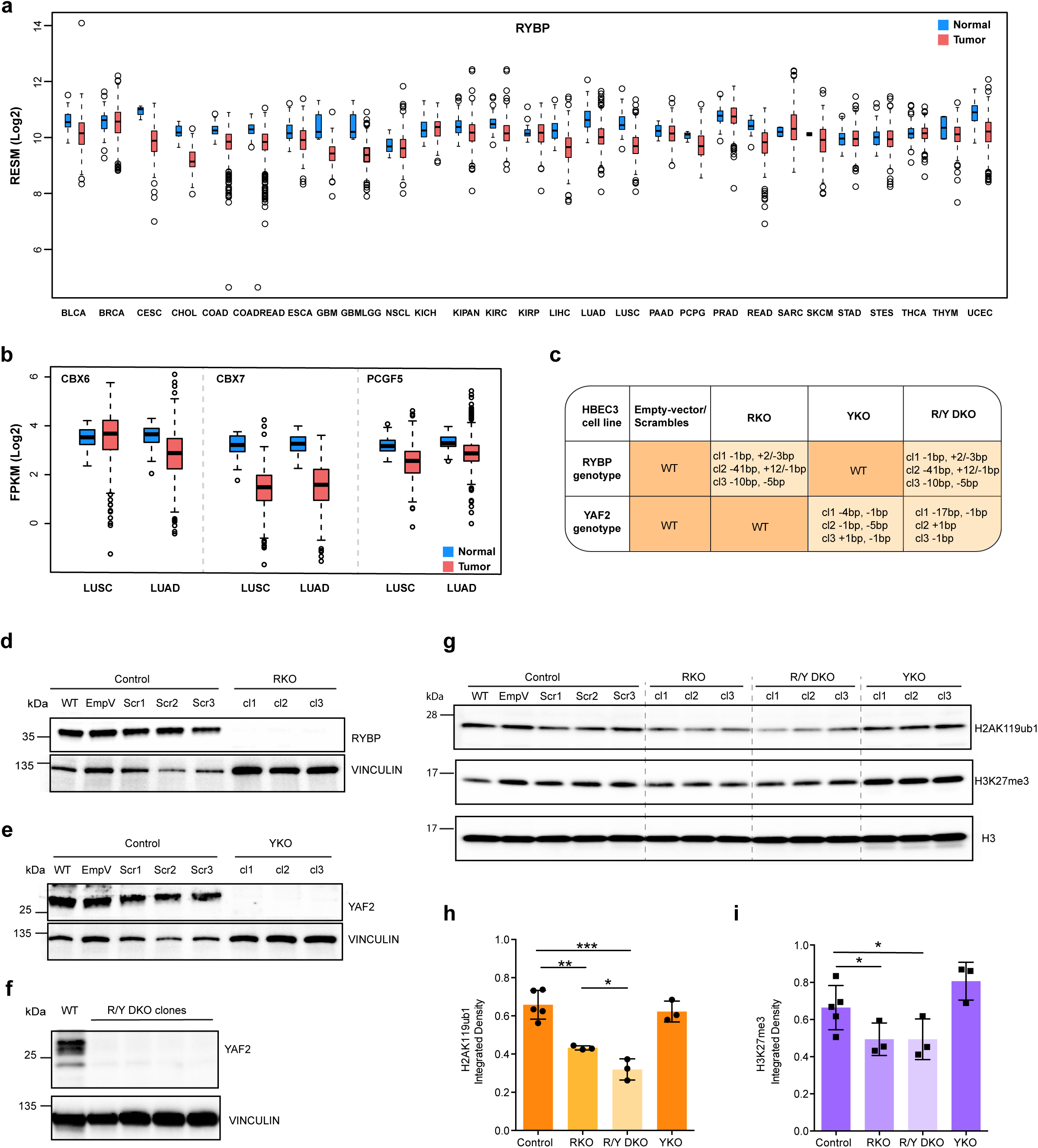
Expression of Polycomb genes in cancer and inactivation of RYBP and YAF2 in HBEC3 cells. **a.** mRNA expression of *RYBP* in various human tumor samples in the TCGA dataset. **b.** mRNA expression of *CBX6*, *CBX7* and *PCGF5* in human lung adenocarcinomas (LUAD, 537; normal solid tissue, 59) and lung squamous cell carcinoma (LUSC, 502; normal solid tissue, 49) in the TCGA dataset. **c.** Table of genotypes of RYBP or YAF2 in controls, RKO, YKO, and R/Y DKO cells. The mutations of the three clones are shown. **d, e, f.** Representative immunoblots of RYBP, YAF2 and VINCULIN in control, RKO, YKO, and R/Y DKO cells. **g.** Representative immunoblots of H2AK119ub1, H3K27me3 and H3 in controls, RKO, DKO or YKO cells. **h.** Quantitation of signals of H2AK119ub1, H3K27me3 and H3 immunoblots by Image J, normalized to the H3 signal. Each dot denotes one clone. Data represents mean ± SD, \**p* < 0.05, \*\**p* < 0.01, \*\*\**p* < 0.001.

**Extended Data Fig. 2.**
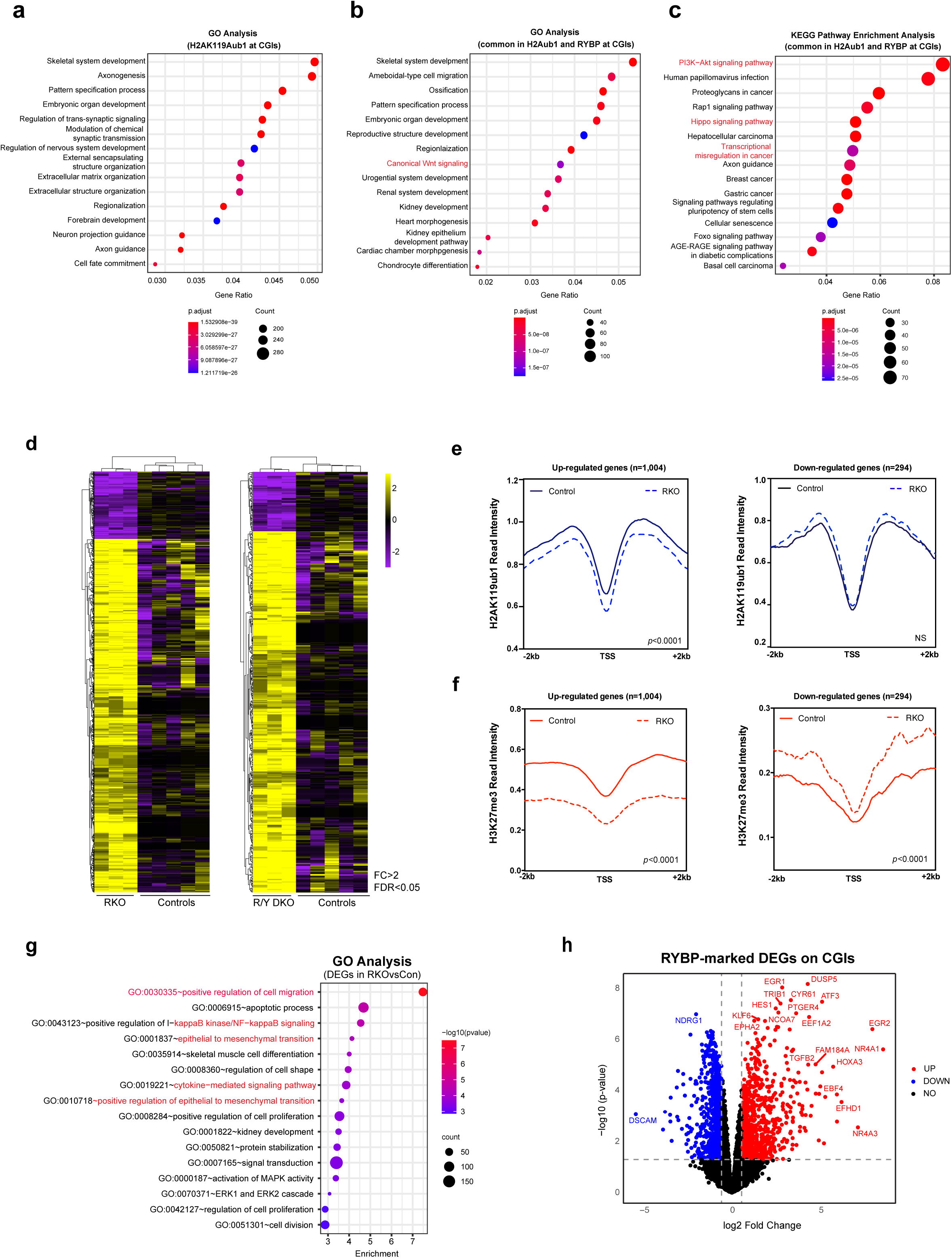
Gene ontology analysis of Polycomb-marked CpG islands and RNA-seq analysis of RYBP knockout cells. **a.** Gene ontology (biological process) enrichment analysis of genes targeted by H2AK119ub1 at their CGIs in R/T123 QKO cells**. b, c.** KEGG pathway and gene ontology (biological process) enrichment analysis of genes targeted by both H2AK119ub1 and RYBP at their CGIs in R/T123 QKO cells. The combined analysis of three independent clones is presented in panels a-c. Some cancer-related pathways are shown in red. **d.** Heatmap of RNA-seq of controls, RKO, and R/Y DKO cells. Each row represents the expression level of a differentially expressed gene ranked by log2- fold change in RKO or R/Y DKO. Five independent clones in control cell lines and three independent clones in different knockout cell lines are presented. The combined analysis is presented. **e, f.** Intensity plot illustrating H2AK119ub1 and H3K27me3 occupancy in controls and RKO cells around the TSS ± 2kb of up-regulated genes and down-regulated genes, respectively. The combined analysis of three independent clones is presented. *P* value is indicated. NS represents no significant difference. **g.** Gene ontology (biological process) enrichment analysis of differentially expressed genes in RKO cells. The combined analysis of three independent clones is presented. Some cancer-related pathways are shown in red. **h.** Volcano plots showing differentially expressed genes (RKO versus controls) with RYBP-marked CGIs. Dotted lines represent the fold change and *p* value cutoffs for significantly enriched genes.

**Extended Data Fig. 3.**
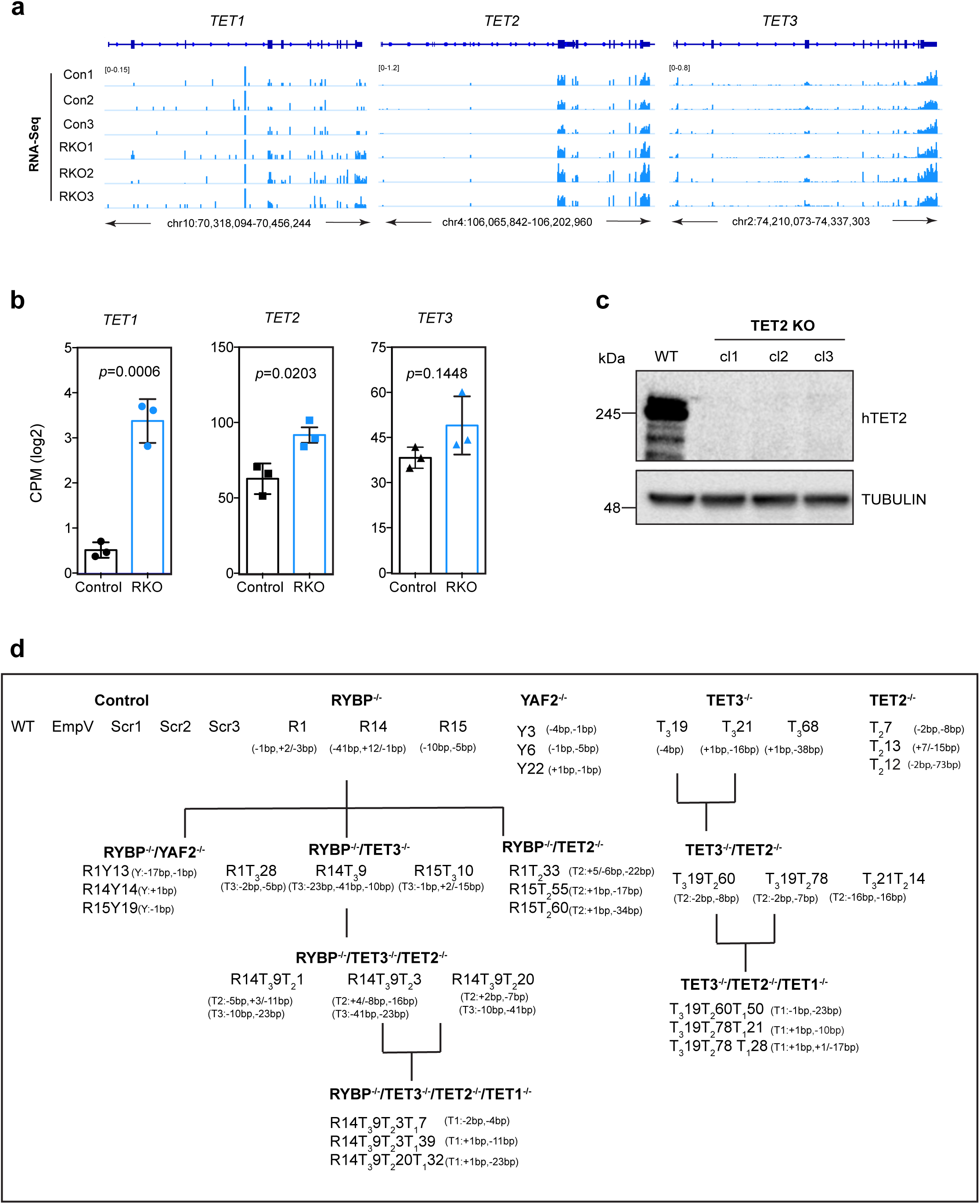
Expression levels of TET genes and Clone Tree. **a.** IGV browser views of the *TET1*, *TET2* and *TET3* loci for RNA-seq data acquired from controls and RKO cells. **b.** RNA-seq data, shown in FPKM values for *TET1*, *TET2* and *TET3* in controls and RKO. Three independent clones are presented. Data represents mean ± SD, *P* values are indicated. **c.** Representatives immunoblot of TET2 and TUBULIN in wildtype and three independent TET2 KO cells. We did not identify antibodies that work for human TET1 or TET3. **d.** Evolutionary tree of knockout clone generation. Specific mutations of *RYBP*, *YAF2*, *TET1*, *TET2* and *TET3* in the different defective clone cells are indicated. Three independent clones for each knockout were obtained.

**Extended Data Fig. 4.**
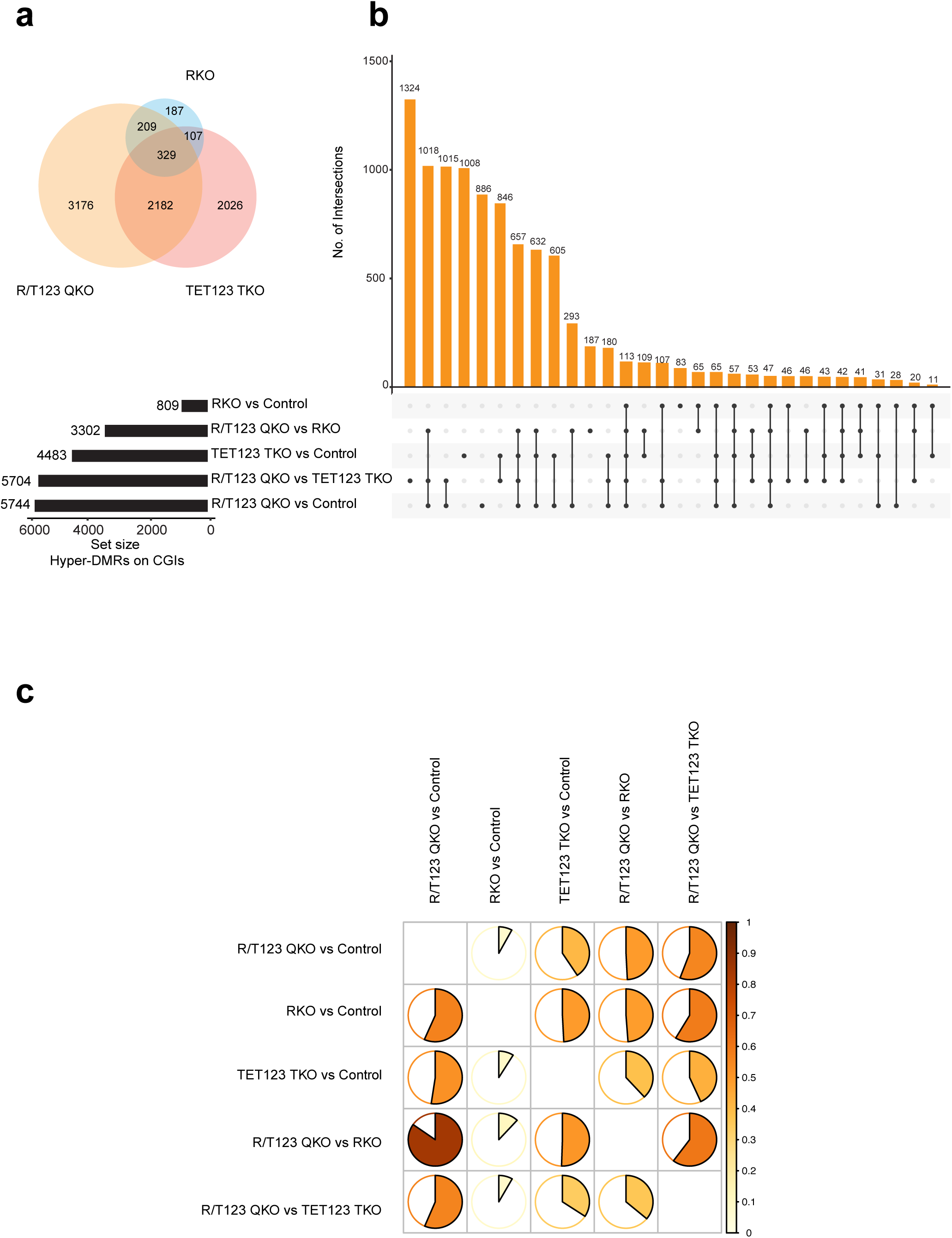
The intersection of DMRs at CGIs. **a.** Venn diagram of overlap of hypermethylated CpG islands between RYBP single knockout (RKO), TET TKO and RYBP/TET1/2/3 QKO. **b.** Upset plot indicating the overlap of DMRs at CGIs from the groups that are indicated. The number on the left side of the black bar indicates the number of the hypermethylated regions at CGIs. **c.** Pairwise plot showing the overlap of percentage of DMRs from the groups that are indicated. Color bars indicate the percentage of hypermethylated CGI overlap from 0% to 100%.

**Extended Data Fig. 5.**
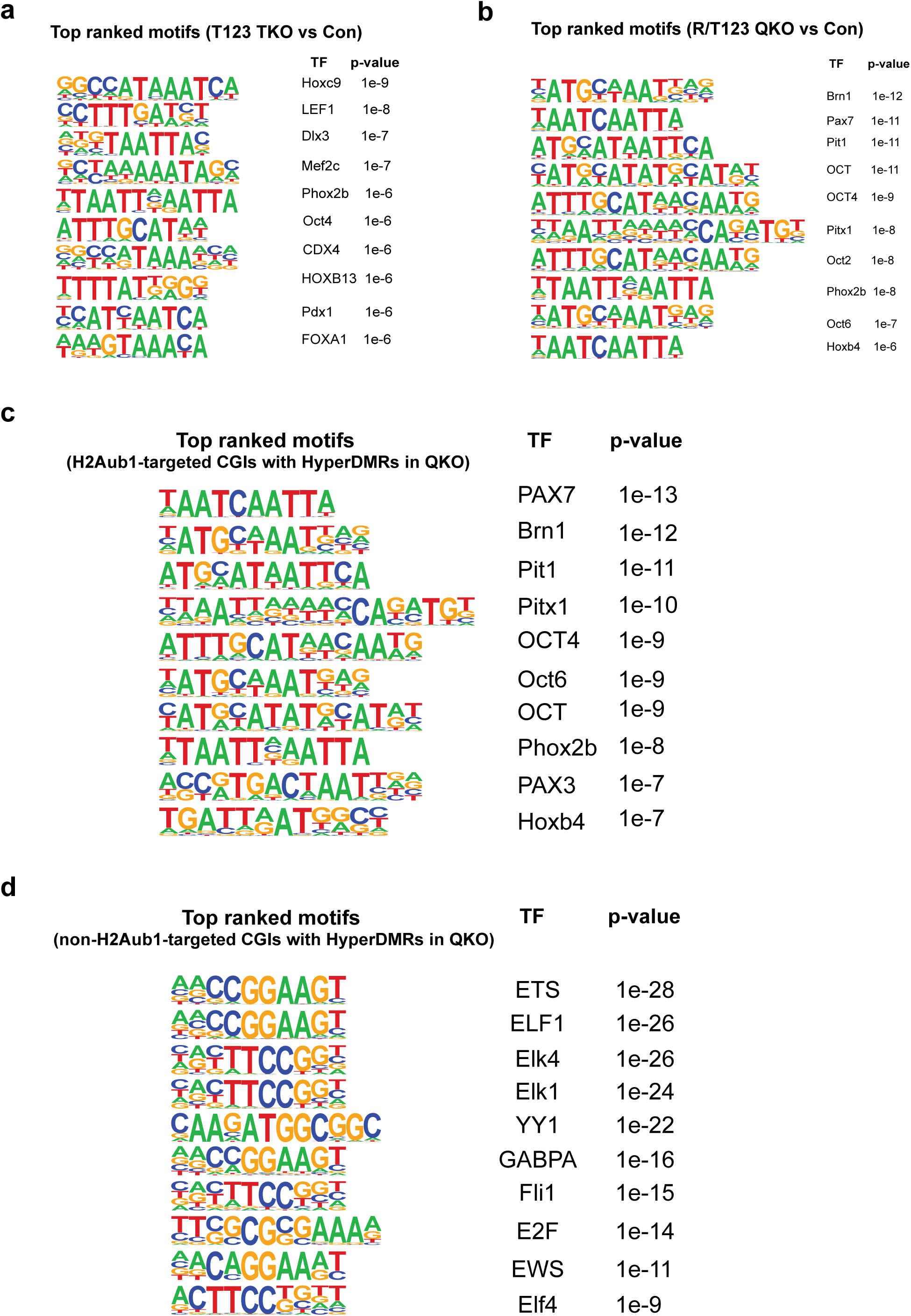
Enrichment of transcription factor motifs in the hypermethylated regions. **a, b.** The top ten enriched DNA binding motifs identified by HOMER known motif analysis in the hypermethylated regions in the TET123 TKO and R/T123 QKO, respectively. **c.** The top ten enriched DNA binding motifs identified by HOMER known motif analysis in the hypermethylated- CGIs that are marked by H2AK119ub1 in the R/T123 QKO cells. **d.** The top ten enriched DNA binding motifs identified by HOMER known motif analysis in the hypermethylated-CGIs that are *not* marked by H2AK119ub1 in the R/T123 QKO cells.

**Extended Data Fig. 6.**
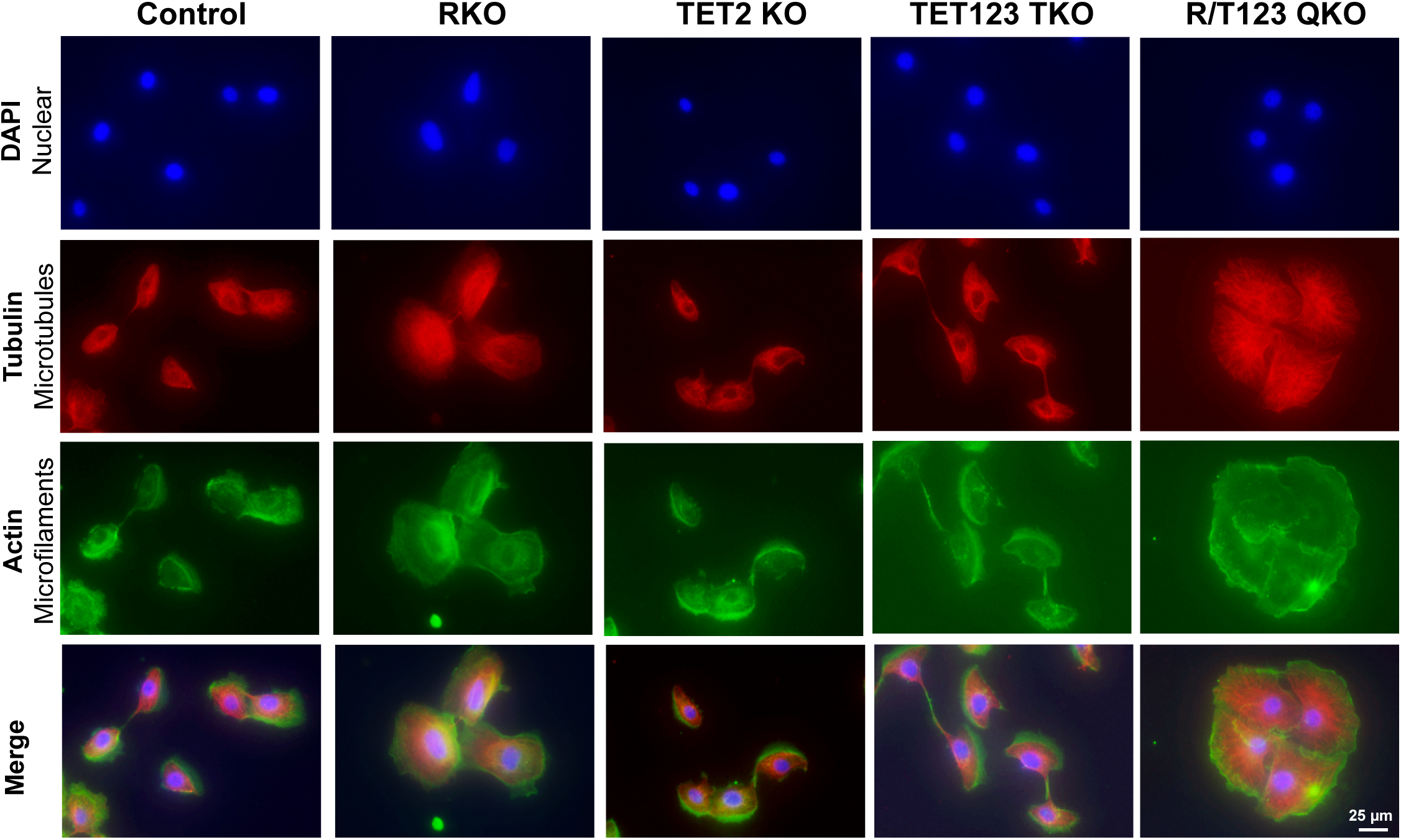
Cell morphology and cytoskeleton in the RYBP and TET defective cells. Immunofluorescence staining of actin (microfilaments) and tubulin (microtubules) in controls, RKO, TET2 KO, TET123 TKO, and R/T123 QKO cells. Tubulin is displayed in red, actin in green, DAPI-stained nuclei in blue. Magnification is the same in all pictures, scale bar, 25 μm.

**Extended Data Fig. 7.**
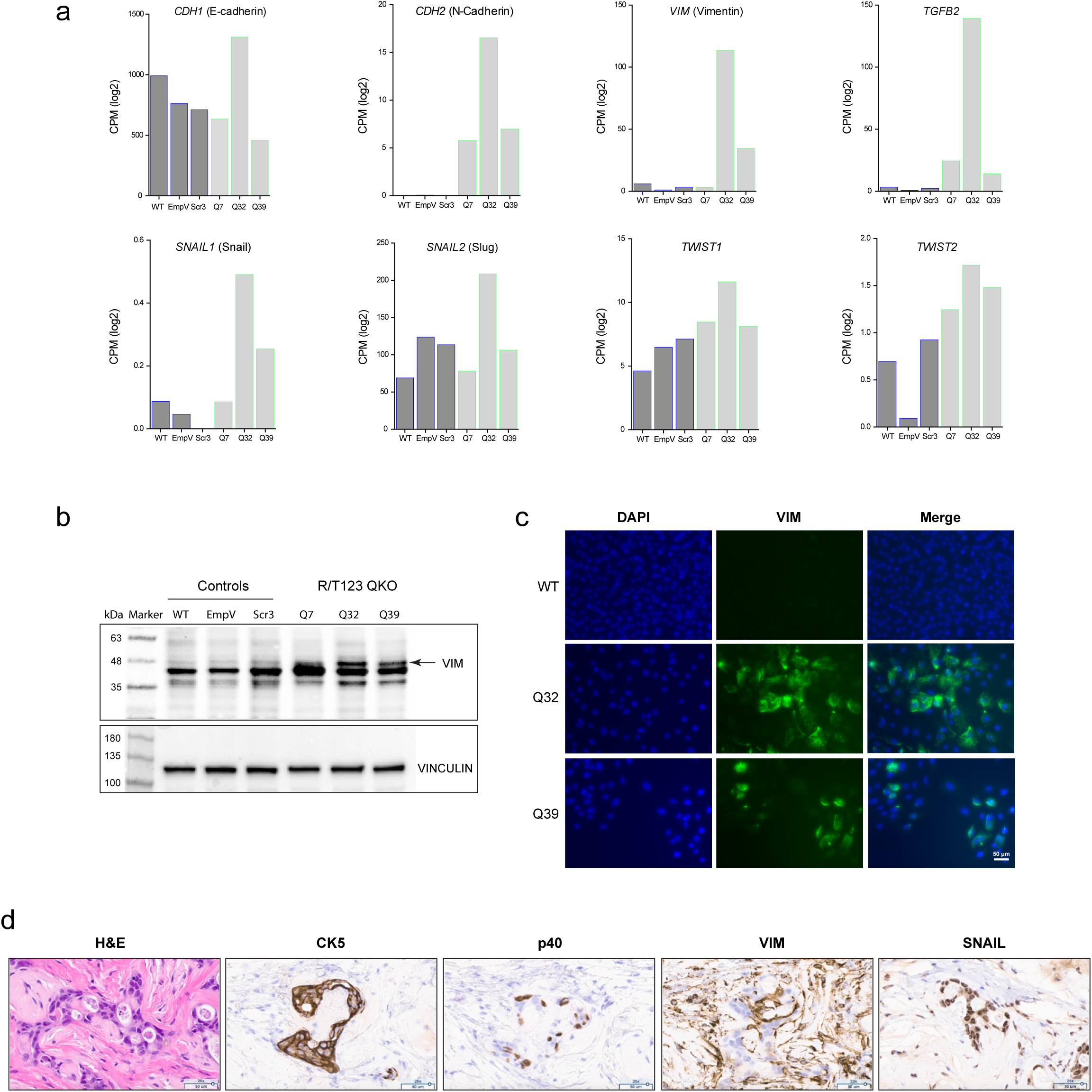
Epithelial to mesenchymal transition in the Q32 QKO cells. **a.** RNA-seq data, shown in FPKM values for the EMT-relevant genes *CDH1*, *CDH2*, *VIM*, *SNAIL1*, *SNAIL2*, *TWIST1*, *TWIST2*, and *TGFB2* in controls and R/T123 QKO cells. Three independent clones are presented. **b.** Western blot with anti-vimentin antibody in controls and QKO cells. **c.** Immunofluorescence staining. VIM is displayed in green, DAPI-stained nuclei in blue. Magnification is the same in all pictures; scale bar, 50 μm. **d.** Representative histology of xenografts obtained from mice injected with R/T123 QKO cell (clone QKO32). Adjacent sections were stained with different antibodies. Representative sections were stained with H&E, CK5, p40, VIM and SNAIL (SNAI1/2) antibodies. Scale bars, 50 µm.

**Extended Data Fig. 8.**
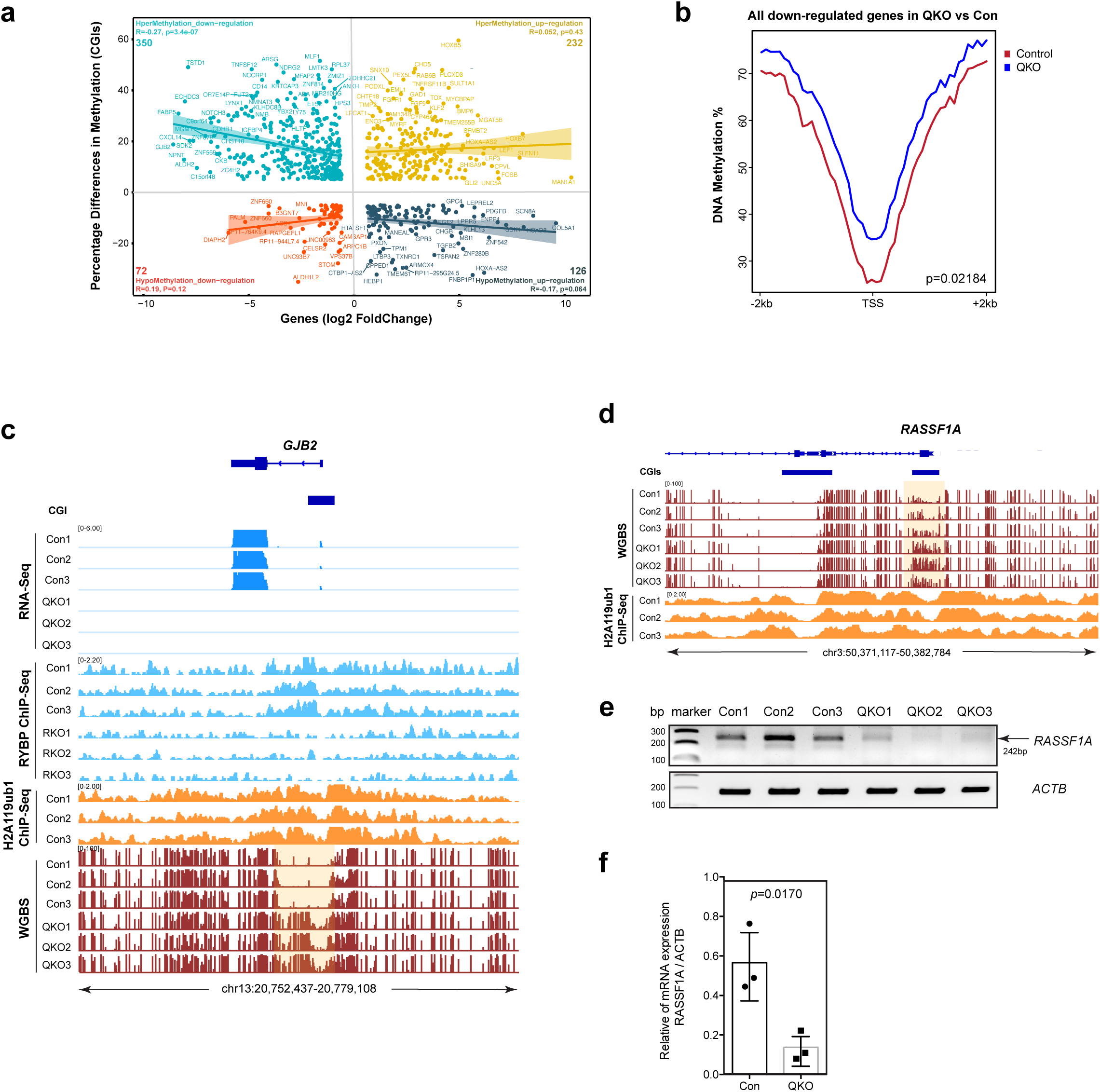
Methylation and gene expression analysis in the QKO. **a.** Correlation between the DNA methylation changes of DMRs at CGIs and expression of DMR- associated genes in R/T123 QKO cells. **b.** Metaplot illustrating DNA methylation levels around the TSS ± 2kb of down-regulated genes in controls and R/T123 QKO cells. *P* value is indicated. The combined analysis of three independent clones is presented in (a,b) **c.** Hypermethylation and downregulation of the *GJB2* gene. RNA-seq, RYBP ChIP-seq, H2AK119ub1 ChIP-seq, and WGBS data are shown for the *GJB2* gene locus in controls, RKO or R/T123 QKO cells. Light orange shading indicates the DMR at the promoter-associated CGI. Three independent clones are presented. **d-f.** Hypermethylation and downregulation of the *RASSF1A* gene. **d.** WGBS and H2AK119ub1 data are shown for the promoter and CpG island of the *RASSF1A* isoform. Light orange shading indicates the DMR. **e.** RT-PCR analysis of *RASSF1A* and *ACTB* control. **f.** Quantitation of *RASSF1A* expression in three independent clones. The P value is indicated.

**Extended Data Fig. 9.**
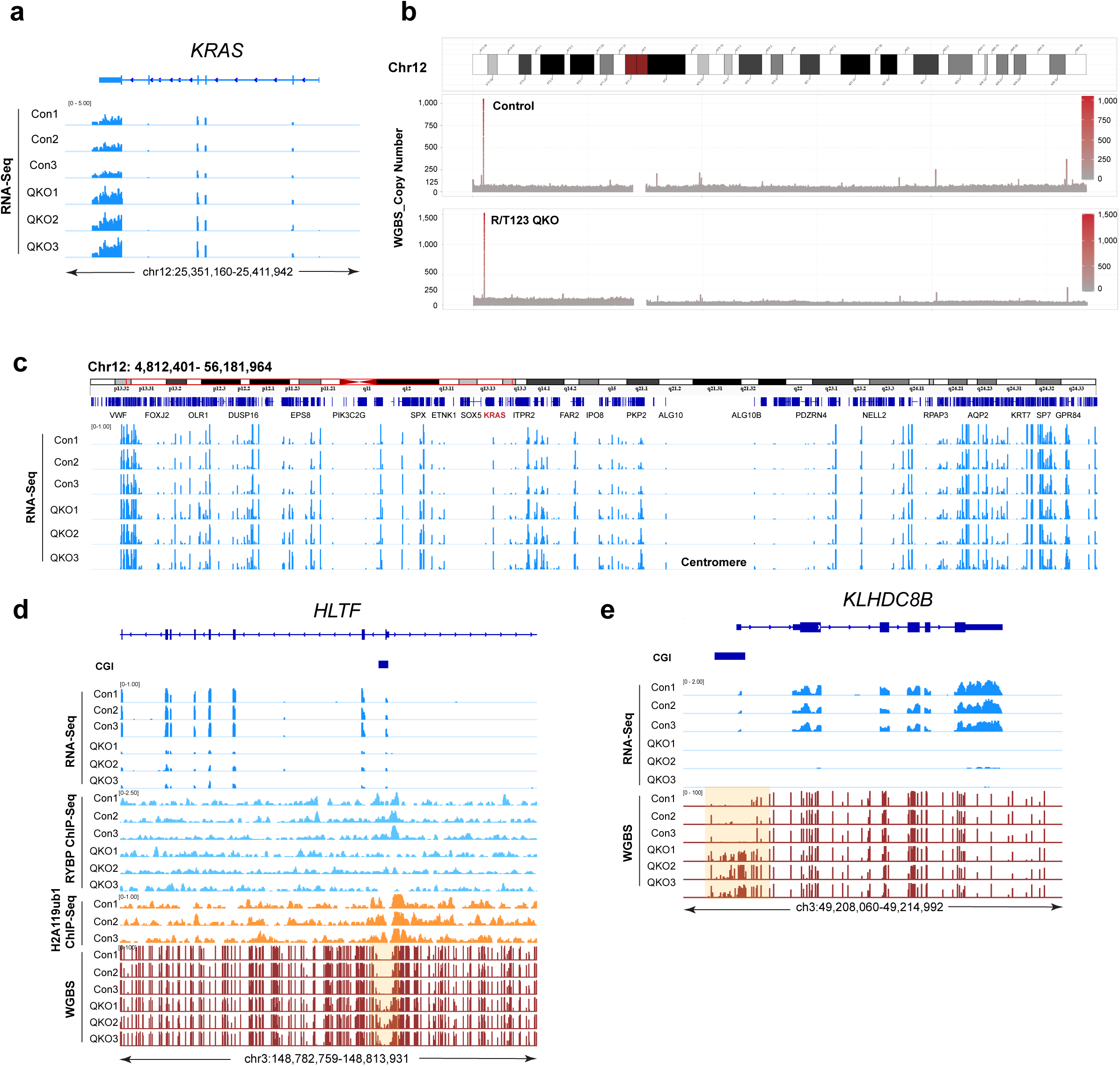
Genomic instability after inactivation of RYBP and TET proteins. **a.** IGV browser views of the *KRAS* locus for RNA-seq data acquired from controls and R/T123 QKO cells. **b.** Copy number at chromosomes 12 from WGBS data from controls and R/T123 QKO cells. **c.** IGV browser views of the full short arm and partial long arm of chromosome 12 for RNA-seq data acquired from controls and R/T123 QKO cells. The centromere is indicated. **d.** RNA-seq, RYBP ChIP-seq, H2AK119ub1 ChIP-seq, and WGBS at the *HLTF* gene locus and its CGI in controls, RKO or R/T123 QKO cells. Light orange shading indicates DMRs at the CGI. **e.** RNA-seq and WGBS at the *KLHDC8B* gene locus in controls and R/T123 QKO cells. Light orange shading indicates the DMR at the promoter-associated CGI. Three independent clones are presented at (a-e).

**Extended Data Fig. 10.**
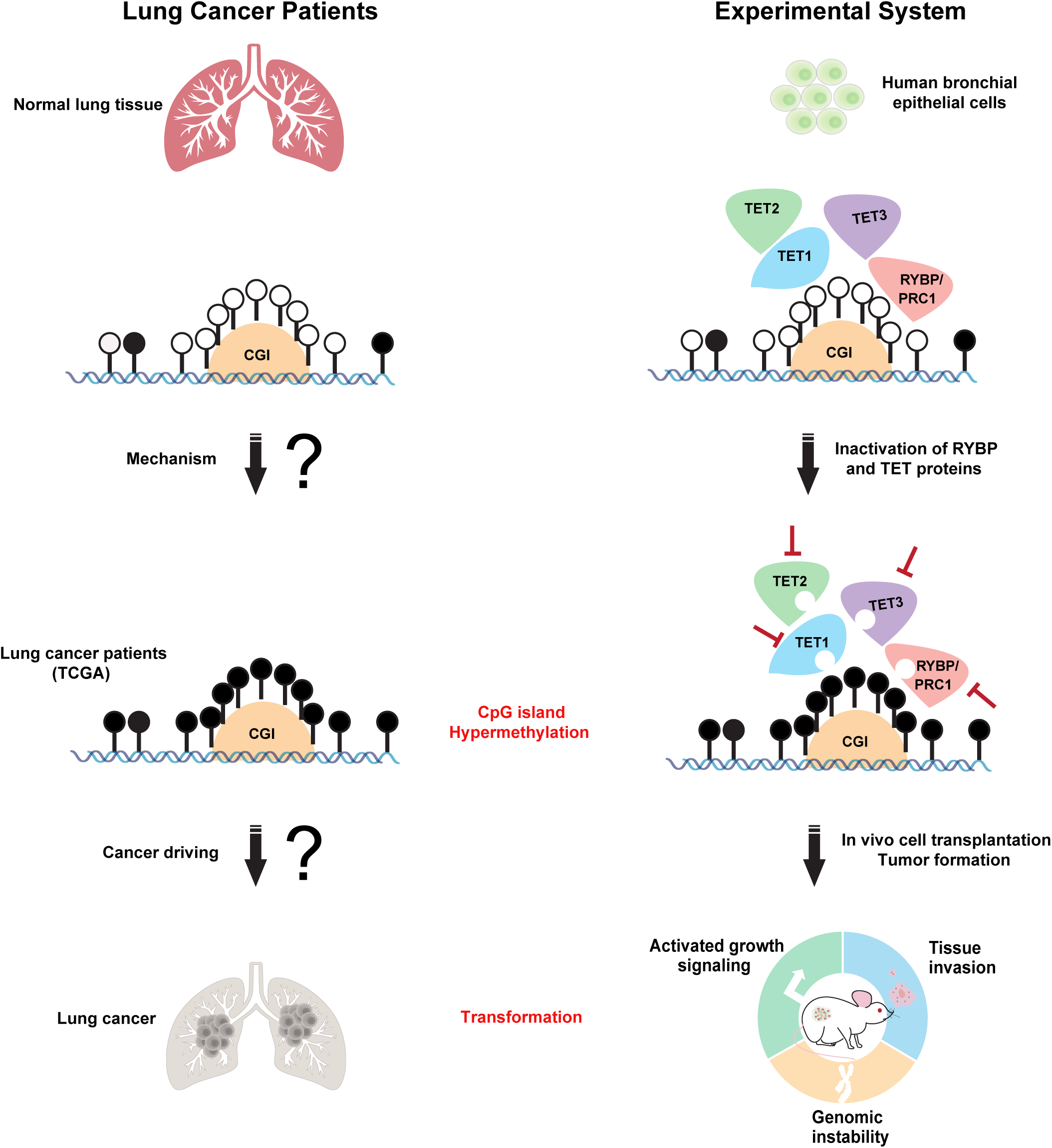
Model of DNA hypermethylation in cancer and lung tumor formation. The left side of the diagram shows the DNA hypermethylation events observed in patients with lung squamous cell carcinomas and the open questions (mechanisms? cancer driving events?). The rights side displays our experimental system of bronchial epithelial cells with dysfunction of RYBP and TET proteins leading to CpG island hypermethylation with the consequences of cell transformation and tumorigenesis.

## Notes

### Competing Interest Statement

The authors have declared no competing interest.

