## Supplemental Tables for "Dysfunction of the Polycomb protein RYBP and of 5-methylcytosine oxidases leads to widespread CpG island hypermethylation and cell transformation": Supplementary Table 1.docx

**Supplementary Table 1**. List of HBEC3 clonal KO cell lines. Mutated DNA sequences are indicated.

| **Cell line** | **Genes** | **Clone#** | **Mutant sequences:**  **(N) denotes insertion, -- denotes deletion** | **Genotype** |
| --- | --- | --- | --- | --- |
| RYBP KO  (RKO) | RYBP | cl1 | AAAACCTCG - ATCAATTCTCAGCTGGTGGCACAACAAGT  AAAACC (CC) --- ATCAATTCTCAGCTGGTGGCACAACAAGT | -1bp  +2/-3bp |
|  |  | cl14 | AAAACCTCGG (CTGGTTTCAATT) - TCAATTCTCAGCTGGTGGCACAAC  -----------------------------------------TCTCAGCTGGTGGCACAAC | +12/-1bp  -41bp |
|  |  | cl15 | AAAACCTC ----- AATTCTCAGCTGGTGGCACAACAAGT  AAAACCTC ---------- TCAGCTGGTGGCACAACAAGT | -5bp  -10bp |
| YAF2 KO  (YKO) | YAF2 | cl3 | GGGACTGTAGCGTCTGCACC ---- GGAACAGCGCCG  GGGACTGTAGCGTCTGCAC - TTCCGGAACAGCGCCG | -4bp  -1bp |
|  |  | cl6 | GGGACTGTAGCGTCTGCAC - TTCCGGAACAGCGCCG  GGGACTGTAGCGTC ----- CTTCCGGAACAGCGCCG | -1bp  -5bp |
|  |  | cl22 | GGGACTGTAGCGTCTGCACC (C) TTCCGGAACAGCGCCG  GGGACTGTAGCGTCTGCAC - TTCCGGAACAGCGCCG | +1bp  -1bp |
| RYBP/YAF2 DKO  (R/Y DKO) | YAF2 | cl13 | GGGACTGTAGCGTCTGC ----------------- CG GGGACTGTAGCGTCTGCAC - TTCCGGAACAGCGCCG | -17bp  -1bp |
|  |  | cl14 | TCAAGTGCATGATGTGCGAT (T) GTGCGGAAG  TCAAGTGCATGATGTGCGAT (T) GTGCGGAAG | +1bp  +1bp |
|  |  | cl19 | GGGACTGTAGCGTCTGCACCTTC - GGAACAGCGCCG  GGGACTGTAGCGTCTGCACCTTC - GGAACAGCGCCG | -1bp  -1bp |
| TET2 KO  (T2 KO) | TET2 | cl7 | TCACG -------- TTATTTGACCATAAGGCTCTTACTCTCAAATCA  TCACGCCAA -- CGTTATTTGACCATAAGGCTCTTACTCTCAAATCA | -8bp  -2bp |
|  |  | cl12 | TCACGCCAA -- CGTTATTTGACCATAAGGCTCTTACTCTCAAATCA  -------------------------------------------------------------------------CA | -2bp  -73bp |
|  |  | cl13 | TCACGCCAAG (AAACTTC)--------------- TCGTTATTTGACCAT  TCACGCCAAG (AAACTTC)--------------- TCGTTATTTGACCAT | +7/-15bp  +7/-15bp |
| TET1/TET2/TET3  TKO  (TET123 TKO) | TET1 | cl21 | ACAAAGGCCCATATTATACACA (A) CCTTGGGGCAGGA  ACAAAGGCCCATATT ---------- TGGGGCAGGA | +1bp  -10bp |
|  |  | cl28 | ACAAAGGCCCATATTATACACA (A) CCTTGGGGCAGGA  ACAAAGGCCCATA (C)----------------- CAGGA | +1bp  +1/-17bp |
|  |  | cl50 | ACAAAGGCCCATATTATACACAC - TTGGGGCAGGA  ACAAAGGC --------------------- AGGA | -1bp  -23bp |
|  | TET2 | cl14 | AGTTTCAC ---------------- ACCATAAGGCTCTTACTCTCAAATCACA  AGTTT ---------------- TTGACCATAAGGCTCTTACTCTCAAATCACA | -16bp  -16bp |
|  |  | cl60 | AGTTTCACGCCAAGTC -- TATTTGACCATAAGGCTCTTACTCT  AGTTTCACGCCAAGT -------- GACCATAAGGCTCTTACTCT | -2bp  -8bp |
|  |  | cl78 | AGTTTCACGCCAAGTC -- TATTTGACCATAAGGCTCTTACTCT  AGTTTCAC ------- CGTTATTTGACCATAAGGCTCTTACTCT | -2bp  -7bp |
|  | TET3 | cl19 | TATGGAGAGAAGGGGAAAGCCATCCGGA ---- GAAGGTCAT  TATGGAGAGAAGGGGAAAGCCATCCGGA ---- GAAGGTCAT | -4bp  -4bp |
| TET3 KO  (T3 KO) | TET3 | cl19 | TATGGAGAGAAGGGGAAAGCCATCCGGA ---- GAAGGTCAT  TATGGAGAGAAGGGGAAAGCCATCCGGA ---- GAAGGTCAT | -4bp  -4bp |
|  |  | cl21 | TATGGAGAGAAGGGGAAAGCCATCCGGATC (C) GAGAAGGTCAT  TATGGAGAGAAGGGGA ---------------- GAAGGTCAT | +1bp  -16bp |
|  |  | cl68 | TATGGAGAGAAGGGGAAAGCCATCCGGAT - GAGAAGGTCATCT  TAT -------------------------------------- CT | -1bp  -38bp |
| RYBP/  TET1/TET2/TET3 QKO  (R/T123 QKO) | RYBP | cl14 | AAAACCTCGG (CTGGTTTCAATT) - TCAATTCTCAGCTGGTGGCACA  -----------------------------------------TCTCAGCTGGTGGCACA | +12/-1bp  -41bp |
|  | TET3 | cl9_1 | AAGGGGAAAGCCAT ----------------------- GGGGAAGGA  AAGGGGAAAGCCATCCGG --------- TCATCTACACGGGGAAGGA | -23bp  -10bp |
|  |  | cl9_2 | AAGGGGAAAGCCAT ----------------------- GGGGAAGGA  AAGGG ---------------------------------------- A | -23bp  -41bp |
|  |  | cl9_3 | AAGGGGAAAGCCATCCGG --------- TCATCTACACGGGGAAGGA  AAGGG ---------------------------------------- A | -10bp  -41bp |
|  | TET2 | cl3 | -----------------------------------------TCTCAGCTGGTGGCAC  AAAACCTCGG (CTGGTTTCAATT) - TCAATTCTCAGCTGGTGGCAC | -41bp  +12/-1bp |
|  |  | cl20 | AGTTTCACGCCAAGTCG (CG) TTATTTGACCATAAGGCTCTTAC  AGTTTCACGCCAAGT ------- TGACCATAAGGCTCTTAC | +2bp  -7bp |
|  | TET1 | cl7 | ACAAAGGCCCATATTATAC ---- CTTGGGGCAGGA  ACAAAGGCCCATATTATACACA TT-- GGGGCAGGA | -4bp  -2bp |
|  |  | cl32 | ACAAAGGCCCATATT ---------- GGGGCAGGA  ACAAAGGCCCATATTATACACACC (C)TTGGGGCAGGA | -11bp  +1bp |
|  |  | cl39 | ACAAAGGC ----------------------- AGGA  ACAAAGGCCCATATTATACACA (A) CCTTGGGGCAGGA | -23bp  +1bp |
| TET2/TET3 DKO  (TET2/3 DKO) | TET2 | cl14 | AGTTTCAC ---------------- ACCATAAGGCTCTTACTCTCAAATCACA  AGTTT ---------------- TTGACCATAAGGCTCTTACTCTCAAATCACA | -16bp  -16bp |
|  |  | cl60 | AGTTTCACGCCAAGTC -- TATTTGACCATAAGGCTCTTACTCT  AGTTTCACGCCAAGT -------- GACCATAAGGCTCTTACTCT | -2bp  -8bp |
|  |  | cl78 | AGTTTCACGCCAAGTC -- TATTTGACCATAAGGCTCTTACTCT  AGTTTCAC ------- CGTTATTTGACCATAAGGCTCTTACTCT | -2bp  -7bp |
|  | TET3 | cl19 | TGGAGAGAAGGGGAAAGCCATCCGGA ---- GAAGGTCATCTACA  TGGAGAGAAGGGGAAAGCCATCCGGA ---- GAAGGTCATCTACA | -4bp  -4bp |
|  |  | cl21 | TATGGAGAGAAGGGGAAAGCCATCCGGATC (C) GAGAAGGTCAT  TATGGAGAGAAGGGGA ---------------- GAAGGTCATCT | +1bp  -16bp |
