## Supplemental Tables for "Dysfunction of the Polycomb protein RYBP and of 5-methylcytosine oxidases leads to widespread CpG island hypermethylation and cell transformation": Supplementary Table 7.docx

List of DNA oligonucleotides. List of gRNA Target Sequences. Guide RNAs were used for targeting of the *RYBP*, *YAF2*, *TET1*, *TET2* and *TET3* genes. Listed are the oligonucleotides used for genotyping to identify knockouts of *RYBP*, *YAF2*, *TET1*, *TET2* and *TET3*, and PCR primers for amplifying *RASSF1A*.

| **Gene Name** | **Sequence** |
| --- | --- |
| RYBP gRNA | GCCTAGTTAAGAGTCGACCA |
| YAF2 gRNA#1 | TGTAGCGTCTGCACCTTC |
| YAF2 gRNA#2 | TCACGTACTACACGCTACAC |
| TET1 gRNA | GGCCCATATTATACACACCT |
| TET2 gRNA | TCAGCAATAAACTGGTATTC |
| TET3 gRNA | GATCGAGAAGGTCATCTACA |
| RYBP_ Tclone _F | CAGTAATAAAATACCCAAGGAATGT |
| RYBP_ Tclone _R | GGTGGTGGGGTGGCATA |
| YAF2_ Tclone _F | GGGTCGCAGAATTGGGG |
| YAF2_ Tclone _R | CAGTGTCATGGGAGGGGG |
| TET1_ Tclone _F | GTTTCCTGCGCCCTTTCAT |
| TET1_ Tclone _R | GCTCCCTTGGTTGTCTTTCGT |
| TET2_ Tclone _F | ACAGCACCACCAGAAAACAAAA |
| TET2_ Tclone _R | GCTGGGGTGTGGCTATCAAGT |
| TET3_ Tclone _F | ACCAGAAACTCAAACAAAAGCACAC |
| TET3_ Tclone _R | TGGGGGAAATGACCCTAACC |
| *RASSF1A*_exon4 | GATGAAGCCTGTGTAAGAACCGTCCT |
| *RASSF1A*_exon2αβ | CAGATTGCAAGTTVACCTGCCACTA |
| *ACTB*_F | CCCTGGACTTCGAGCAAGAGAT |
| *ACTB*_R | AAGGTAGTTTCGTGGATGCCACA |
