## Supplemental Tables for "Dysfunction of the Polycomb protein RYBP and of 5-methylcytosine oxidases leads to widespread CpG island hypermethylation and cell transformation": Supplementary Table 8.docx

**Supplementary Table 8**. **List of antibodies.**

Antibody target, suppliers, catalogue number, assay, and amount or dilution used are indicated.

| **Antibody** | **Suppliers** | **Catalogue number** | **Assay** | **Dilution or amount** |
| --- | --- | --- | --- | --- |
| Anti-DEDAF Antibody (RYBP) | Millipore | AB3637 | Western blotting, ChIP-Seq | 1:1000, 2 µg |
| YAF2 | BETHYL | A303-654A-T | Western blotting | 1:1000 |
| VINCULIN | Cell Signaling | 13901 | Western blotting | 1:2000 |
| Histone H3 (D1H2) XP Rabbit | Cell Signaling | 4499 | Western blotting | 1:2000 |
| Ubiquityl-Histone H2A (Lys119) (D27C4) XP Rabbit | Cell Signaling | 8240 | Western blotting, ChIP-Seq | 1:2000, 6 µg |
| Tri-Methyl-Histone H3 (Lys27) (C36B11) | Cell Signaling | 9733 | Western blotting, ChIP-Seq | 1:2000, 2 µg |
| TET2 | Cell Signaling | 18920 | Western blotting | 1:2000 |
| Anti-alpha Tubulin | Abcam | ab7291 | Western blotting, IF | 1:2000 |
| Anti-beta Actin | Abcam | ab8227 | IF | 1:2000 |
| Alexa Fluor® 488 Goat anti-Rabbit | Invitrogen | A11034 | IF | 1:2000 |
| Alexa Fluor® 568 Goat anti-Mouse | Invitrogen | A11004 | IF | 1:1000 |
| Anti-p40 - DeltaNp63 antibody [BC28] | Abcam | ab172731 | IHC | 1:100 |
| Recombinant Anti-Cytokeratin 5 antibody [EP1601Y] - Cytoskeleton Marker | Abcam | ab52635 | IHC | 1:200 |
| Vimentin | Cell Signaling | 5741 | IHC | 1:200 |
| Vimentin | Cell Signaling | 3932 | Western blotting, IF | 1:2000, 1:200 |
| Anti-SNAIL+SLUG antibody | Abcam | Ab85936 | IHC | 1:100 |
